# An Optimized Screen Reduces the Number of GA Transporters and provides Insights into NPF Substrate Determinants

**DOI:** 10.1101/670174

**Authors:** Nikolai Wulff, Heidi Asschenfeldt Ernst, Morten Egevang Jørgensen, Sophie Lambertz, Tobias Maierhofer, Zeinu Mussa Belew, Christoph Crocoll, Mohammed Saddik Motawia, Dietmar Geiger, Flemming Steen Jørgensen, Osman Mirza, Hussam Hassan Nour-Eldin

## Abstract

Based on recent *in vitro* data, a relatively large number of the plant Nitrate transporter 1/Peptide transporter Family (NPF) proteins has been suggested to function as gibberellic acid (GA) transporters. Most GA transporting NPF proteins also appear to transport other structurally unrelated phytohormones or metabolites. Several of the GAs used in previous *in vitro* assays are membrane permeable weak organic acids whose movement across membranes are influenced by the pH-sensitive ion-trap mechanism. Moreover, a large proportion of *in vitro* GA transport activities have been demonstrated indirectly via long-term yeast-based GA-dependent growth assays that are limited to detecting transport of bioactive GAs. Thus, there is a need for an optimized transport assay for identifying and characterizing GA transport. Here, we develop an improved transport assay in *Xenopus laevis* oocytes wherein we directly measure movement of six different GAs across oocyte membranes over short time. We show that membrane permeability of GAs in oocytes can be predicted based on number of oxygen atoms and that several GAs do not diffuse over membranes regardless of changes in pH values. In addition, we show that small changes in internal cellular pH can result in strongly altered distribution of membrane permeable phytohormones. This prompts caution when interpreting heterologous transport activities. We use our transport assay to screen all *Arabidopsis thaliana* NPF proteins for transport activity towards six GAs (two membrane permeable and four non-permeable). The results presented here, significantly reduce the number of *bona fide* NPF GA transporters in *Arabidopsis* and narrow the activity to fewer subclades within the family. Furthermore, to gain first insight into the molecular determinants of substrate specificities towards organic molecules transported in the NPF, we charted all surface exposed amino acid residues in the substrate-binding cavity and correlated them to GA transport. This analysis identified distinct residues within the substrate-binding cavity that are shared between GA transporting NPF proteins; the potential roles of these residues in determining substrate specificity are discussed.

## 1 Introduction

GAs were detected in phloem sap more than 50 years ago suggesting that GAs are mobile phytohormones (Hoad and Bowen, 1968). This has since been supported by multiple studies indicating that GAs can move long distances *in planta* and that transport processes generate local concentration maxima that may be essential for the regulatory roles of GA (Regnault et al., 2015;Regnault et al., 2016;Binenbaum et al., 2018). However, due to lack of molecular knowledge on GA transport, the physiological role of GA mobility remains unclear.

Within recent years, novel *in vitro* and *in vivo* approaches led to the identification of a large number of putative GA transporters (*>*25 different genes in *Arabidopsis* summarized in (Supplementary Table 1). The majority of these genes belong to the NPF (Kanno et al., 2012;Chiba et al., 2015;Saito et al., 2015;Tal et al., 2016). The physiological roles of two NPF-GA transporters have been investigated *in planta*, namely NPF3.1 and NPF2.10 (Saito et al., 2015;David et al., 2016;Tal et al., 2016). The GA related phenotypes in plants mutated in these genes are limited compared with those observed in GA-deficient or GA signaling mutants (Sun, 2008;Plackett et al., 2011). The relatively large number of potential NPF-GA transporters implies that there may be widespread functional redundancy among these transporters. Accordingly, experiments in which expression of multiple GA transporting NPF members are inhibited or knocked-out simultaneously may be needed to unveil their distinct roles (Binenbaum et al., 2018).

The NPF proteins are plant specific subfamily members of the Proton-coupled Oligopeptide Transporter (POT) family, which exists in all kingdoms of life and whose members are important for transport of di- and tripeptides across membranes in symport with at least one proton. In humans, there are four POT family members, two of which are prime targets for drug delivery owing to their central importance for delivery of peptidomimetic drugs to intestinal epithelial cells (Daniel and Kottra, 2004). Crystal structures of bacterial POT family members have identified key residues in the POT substrate-binding cavity which interact with the peptidomimetic substrates and are located in a large cavity able to accommodate nearly limitless variations in amino acid side chains and conjugated species (Biegel et al., 2006;Doki et al., 2013;Guettou et al., 2014;Lyons et al., 2014;Newstead, 2015;2017). Moreover, a conserved ExxE[K/R] motif plays an essential role in intra-transporter salt bridge formations that enable coupling between proton and substrate transport to ensure active transport (Solcan et al., 2012;Aduri et al., 2015).

In *Arabidopsis*, 53 NPF members exist, which are divided into 8 distinct subclades (Léran et al., 2014). Low affinity nitrate transport activities have been demonstrated in members from subclades 1, 2, 3, 4, 5, 6 and 7; peptide transport in subclades 5 and 8; glucosinolate transport in subclade 2; and transport of the phytohormones abscisic acid (ABA), GA and jasmonoyl-isoleucine (JA-Ile) transport activities have been demonstrated for members from subclades 1, 2, 3, 4, 5 and 8, as recently reviewed (Corratge-Faillie and Lacombe, 2017;Wang et al., 2018). Hence, for some NPF substrates, transport activity appears relatively confined to distinct subclades with high degree of amino acid identity (e.g. the glucosinolate transporters GTR1-3 (NPF2.9-NPF2.11) with >60% identity and the peptide transporters (NPF8.1-8.3) with > 55% identity - excluding PTR3 (NPF5.2). In contrast, for other substrates such as the 25 putative NPF-GA transporters and the many low-affinity nitrate transporters from *Arabidopsis* (Supplementary Table 1); there is no apparent phylogenetic clustering within distinct subclades. This discrepancy could be due to features other than determinants of GA substrate specificity weighing higher towards defining phylogenetic differentiation. Alternatively, it is conceivable that the current number of putative GA transporters may be overestimated. The ExxE[K/R] motif plays a role in coupling proton and substrate transport in the NPF proteins but otherwise our knowledge on the transport mechanism of NPF members remains limited (Jørgensen et al., 2015). Analysis of the recent crystal structures of *Arabidopsis* NPF6.3 suggested that its enigmatic dual affinity transport mode is controlled by Thr101 phosphorylation-dependent dimerization (Liu and Tsay, 2003) and that nitrate binds to His356 that is only conserved in one out of the ∼20 other suggested NPF nitrate transporters (Parker and Newstead, 2014;Sun et al., 2014;He et al., 2017;Wang et al., 2018). The structure of NPF6.3 provides a welcomed basis for inferring structure-function relationships for the NPF proteins but has so far not been used to explain the molecular determinants of substrate specificities of any NPF transporter towards organic molecules.

The knowledge gap in our understanding of the relationship between structure and function of NPF transporters complicates the identification NPF transporter substrates based on phylogenetic relationships. Hence, understanding the intricate details of substrate selectivity among plant NPF members will be crucial for predicting and elucidating physiological functions.

Here, we aim to increase our understanding of the relationship between structure and function of plant NPF-GA transporters by searching for distinct features among potential substrate-interacting residues within the substrate-binding cavity of GA transporting NPF proteins. In this context, it is paramount to ascertain the *in vitro* data assigning GA transport activity to individual NPF members. To achieve this, we develop and optimize a short-term quantitative GA transport assay in *Xenopus* oocytes that in theory is capable of identifying both GA importers and exporters, and simultaneously detects transport of intermediate-, bioactive- and catabolic GA species. We use the optimized transport assay to identify candidate *Arabidopsis* NPF-GA transporters by characterizing and critically assessing the GA substrate preference map among all *Arabidopsis* NPF members. A surprising result from our study is that some NPF proteins are capable of profoundly altering intracellular pH in *Xenopus* oocytes. In a case study, we show that such alterations may lead to changed accumulation equilibria of membrane permeable phytohormones. In addition, we generate a map of all amino acid residues within the substrate-binding cavity of NPF proteins that may play a role in determining substrate specificity and use the GA substrate preference map to correlate GA transport activity to structural features of NPF proteins. Our findings provide a set of critical considerations that will help in guiding physiological investigations.

## 2 Material and Methods

### 2.1 Cloning the entire NPF and cRNA generation

Coding DNA sequences of the entire *Arabidopsis* NPF were cloned into the *Xenopus* oocyte expression vector, pNB1u as indicated in (Supplementary Table 2) (Nour-Eldin et al., 2006). 17 of the CDS were obtained as uracil-containing non-clonal DNA fragments, codon optimized for *Xenopus* oocyte expression from Thermo Fisher Scientific Geneart wherein a uracil had been placed at either end to facilitate direct insertion into pNB1u via USER cloning (Jørgensen, 2017). Nine of the CDS were amplified via PCR from *Arabidopsis* cDNA. 15 of the CDS were amplified from vectors obtained from Pro. Wolf B. Frommer or Dr. Eilon Shani. Nine of the CDS were amplified from vectors obtained from RIKEN Bioresource Center (BRC) (Seki et al., 1998;Seki et al., 2002). The remaining three CDS had been cloned into pNB1u in a previous study (Nour-Eldin et al., 2012). Primers used for PCR amplification are indicated in (Supplementary Table 2). All PCR amplified fragments had a uracil incorporated at either end via the primers and were cloned into pNB1u via USER cloning as described previously (Nour-Eldin et al., 2006). We used the X7 polymerase (Nørholm, 2010) in the PCR reactions. Linearized DNA templates for RNA synthesis were obtained by PCR amplifying the coding sequences surrounded by *Xenopus* β-Globin 5’- and 3’- UTRs from pNB1u using forward primer (5’ – AATTAACCCTCACTAAAGGGTTGTAATACGACTCACTATAGGG – 3’) and reverse primer (5’ – TTTTTTTTTTTTTTTTTTTTTTTTTTTTTATACTCAAGCTAGCCTCGAG – 3’) PCR products were purified using E.Z.N.A Gel extraction kit (Omega Bio-tek) using the manufacturer’s instructions. PCR products were *in vitro* transcribed using the mMessage mMachine T7 transcription kit (InVitrogen) using the manufacturer’s instructions.

### 2.2 Synthesis of JA-Ile

JA-Ile conjugate was chemically synthesized as described in (Kramell et al., 1988).

### 2.3 *Xenopus* oocyte transport assays

Defolliculated *Xenopus laevis* oocytes (stage V-VI) were purchased from Ecocyte Biosciences and were injected with 25 ng cRNA in 50.6 nl (500 ng/µl) using a Drummond Nanoject II and incubated for 2-4 days at 16 °C in HEPES-based kulori (90 mM NaCl, 1 mM KCl, 1 mM MgCl_2_, 1 mM CaCl_2_, 5 mM HEPES pH 7.4) before use. Expressing oocytes were pre-incubated in MES-based kulori (90 mM NaCl, 1 mM KCl, 1 mM MgCl_2_, 1 mM CaCl_2_, 5 mM MES pH 5) for 4 min, before being transferred to phytohormone-containing MES-based kulori for 60 min. Afterwards, oocytes were washed 3 times in 25 ml HEPES-based kulori followed by one wash in 25 ml deionized water, homogenized in 50 % methanol and stored for >30 min at −20 °C. Following centrifugation (25000 g for 10 min 4 °C), the supernatant was mixed with deionized water to a final methanol concentration of 20 % and filtered through 0.22 µm (MSGVN2250, Merck Millipore) before analytical LC-MS/MS as described below.

### 2.4 pH stabilization

Expressing oocytes were injected with 50.6 nl 0.5 M MES 50 mM EGTA adjusted to pH 5.7 with 0.5 M TRIS for stabilizing internal oocyte pH to ∼6.25 or 0.5 M TRIS 50 mM EGTA adjusted to 7.7 with 0.5 M MES for stabilizing internal oocyte pH to ∼7.5. Assays were carried out as described above, with the exceptions that sorbitol was included in the MES and the HEPES-based kulori solutions to a final concentration of 50 mM to avoid oocyte swelling, and the timeframe between injection of pH stabilizing buffer to assay was terminated was held to a maximum 30 min.

### 2.5 Metabolite quantification by LC-MS/MS

Compounds in the diluted oocyte extracts were directly analyzed by LC-MS/MS. The analysis was performed with modifications from the method described in (Tal et al., 2016). In brief, chromatography was performed on an Advance UHPLC system (Bruker, Bremen, Germany). Separation was achieved on a Kinetex 1.7u XB-C18 column (100 x 2.1 mm, 1.7 µm, 100 Å, Phenomenex, Torrance, CA, USA) with 0.05% v/v formic acid in water and acetonitrile (with 0.05% formic acid, v/v) as mobile phases A and B, respectively. The elution profile for GAs, glucosinolates and glycylsarcosine was: 0-0.5 min, 2% B; 0.5-1.3 min, 2-30% B; 1.3-2.2 min 30-100% B, 2.2-2.8 min 100%; 2.8-2.9 min 100-2% B and 2.9-4.0 min 2% B. The elution profile for ABA, jasmonic acid (JA), Ja-Ile and oxo-phytodienoic acid (OPDA) was: 0-0.5 min, 2% B; 0.5-1.2 min, 2-30% B; 1.2-2.0 min 30-100% B, 2.0-2.5 min 100%; 2.5-2.6 min 100-2% B and 2.6-4.0 min 2% B. The mobile phase flow rate was 400 µl_*_min^-1^ and column temperature was maintained at 40 °C. The liquid chromatography was coupled to an EVOQ Elite triple quadrupole mass spectrometer (Bruker, Bremen, Germany) equipped with an electrospray ion source operated in positive and negative ionization mode. Instrument parameters were optimized by infusion experiments with pure standards. For analysis of GAs, glucosinolates and glycylsarcosine the ion spray voltage was maintained at +4000 V and −4000 V in positive and negative ionization mode, respectively and heated probe temperature was set to 200 °C with probe gas flow at 50 psi. For ABA, JA, JA-Ile and OPDA the ion spray voltage was maintained at −3300 V in negative ionization mode and heated probe temperature was set to 120 °C with probe gas flow at 40 psi. Remaining settings were identical for all analytical methods with cone temperature set to 350 °C and cone gas to 20 psi. Nebulizing gas was set to 60 psi and collision gas to 1.6 mTorr. Nitrogen was used as probe and nebulizing gas and argon as collision gas. Active exhaust was constantly on. Multiple reaction monitoring was used to monitor analyte parent ion → product ion transitions for all analytes: Multiple reaction monitoring transitions and collision energies were optimized by direct infusion experiments. Detailed values for mass transitions can be found in (Supplemental Table 3). Both Q1 and Q3 quadrupoles were maintained at unit resolution. Bruker MS Workstation software (Version 8.2.1, Bruker, Bremen, Germany) was used for data acquisition and processing. Linearity in ionization efficiencies were verified by analyzing dilution series of standard mixtures. Sinigrin glucosinolate was used as internal standard for normalization but not for quantification. Quantification of all compounds was achieved by external standard curves diluted with the same matrix as the actual samples. All GAs were analyzed together in a single method apart from GA12 which suffered from severe ion suppression when combined with the other GAs. Samples with GA12 were therefore analyzed separately and with separate dilution series for quantification. Similarly, other hormones than GAs (ABA, JA, JA-Ile and OPDA) were analyzed in separate analytical runs. Glycylsarcosine and 4-methylthio-3-butenyl were analyzed in a combined analytical run.

### 2.6 pH measurements of oocyte lumen

pH-electrodes were pulled from borosilicate glass capillaries (KWIK-FIL TW F120-3 with filament) on a vertical puller (Narishige Scientific Instrument Lab), baked for 120 min at 220 °C and silanized for 60 min with dimethyldichlorosilane (Silanization Solution I, Sigma Aldrich). Electrodes were backfilled with a buffer containing 40 mM KH_2_PO_4_, 23 mM NaOH and 150 mM NaCl (pH 7.5). The electrode tip was filled with a proton-selective ionophore cocktail (hydrogen ionophore I cocktail A, Sigma-Aldrich) by dipping the tip into the cocktail. Oocytes, as described above, were placed in freshly made HEPES-based ekulori (2 mM LaCl_3_, 90 mM NaCl, 1 mM KCl, 1 mM MgCl_2_, 1 mM CaCl_2_, 5 mM HEPES pH 7.4) for at least 30 min prior to three-electrode voltage clamp experiments. Before each oocyte a pH calibration curve was made for each oocyte using 100 mM KCl pH 5.5, 100 mM KCl pH 6.5 and 100 mM KCl pH 7.5. Oocytes were clamped at 0 mV and perfused with HEPES-based ekulori pH 7.4, followed by MES-based ekulori (2 mM LaCl_3_, 90 mM NaCl, 1 mM KCl, 1 mM MgCl_2_, 1 mM CaCl_2_, 5 mM MES pH 5) and internal pH response was measured continuously as a function of external pH change.

### 2.7 Multiple sequence alignment and structure guided identification of cavity lining residues

In order to bring the GA transport function into a structural context, NPF protein sequences from the 31 plant genomes and two outgroups as performed by (Léran et al., 2014) were retrieved from Phytozome version 9.1(Goodstein et al., 2012) and annotated with their NPF IDs (Léran et al., 2014). Guided by blastp (Altschul et al., 1990), long sequences comprising either multiple NPF modules, or fusions with other proteins, were trimmed to the size of single NPF proteins. Initially, the eight NPF subclades were treated separately, for practical reasons and to facilitate robust multiple sequence alignments produced by MUSCLE (Edgar, 2004). For each group, the *Arabidopsis* NPF6.3 sequence was included as a structural reference (PDB: 4OH3) (Sun et al., 2014). Sequences that due to inserts or gaps were not consistent with an intact Major Facilitator Superfamily (MFS) fold were discarded. In the end, the trimmed multiple sequence alignments for all eight NPF subclades were combined and re-aligned using MAFFT (Katoh and Standley, 2013) followed by manual adjustments. The final alignment comprises 1585 NPF sequences.

In order to embrace structural variability in the substrate binding site, the selection of a subset of amino acid positions that defines the substrate-binding site was guided using four crystal structures of bacterial POT family members, including two complexes with the peptidomimetic drug alafosfalin with different binding modes, as well as homology models constructed to represent the outward-facing conformation of the transporters. The four bacterial POT structures (PDB: 4IKZ, 4LEP, 2XUT and 4APS) (Newstead et al., 2011;Solcan et al., 2012;Doki et al., 2013;Guettou et al., 2013), all in inward-facing conformations, were superimposed and surface-exposed residues within 8 Å from the alafosfalin in either of the structures were extracted and included in the subset of amino acids that makes up the binding site residues.

Importantly, all known POT structures represent inward-open conformations of the transporter, whereas generally substrate recognition from uptake, will occur when the transporter is in the outward-open orientation. In an attempt to identify residues that line the substrate-binding cavity in the outward-facing conformation, homology models were constructed based on outward-open structures of members of the MFS. The two different outward-open structures of FucP and YajR (PDB: 3O7Q and 3WDO) (Dang et al., 2010;Jiang et al., 2013) were used as templates. Sequence alignments of GkPOT (PDB: 4IKZ) (Doki et al., 2013), FucP and YajR sequences were made with PROMALS3D (Pei et al., 2008) and further refined by hand. Outward-facing homology models of GkPOT were made using MODELLER v9.12 (Sali and Blundell, 1993). Finally, surface-exposed residues within 12 Å from the central cavity (as measured from R36-NH1 and W306-CZ2, respectively) were extracted and included in the binding site residue subset. Our final binding site residue subset, defining the substrate-binding cavity, comprises 51 positions. The sequence logos of GA transporters and non-GA transporters were prepared by the WebLogo program (Crooks et al., 2004).

### 2.8 Principal Component analysis

The Principal Component Analysis (PCA) was performed with KNIME using the PCA Compute node (Berthold et al., 2008).

## 3 Results

### 3.1 *Xenopus* oocyte membrane permeability towards GAs at different external pH

GAs are weak organic acids with lipophilic properties. Consequently, when heterologous transport assays utilize external media with an acidic pH, membrane permeable GAs will be subject to the so-called ion-trap mechanism (Rubery and Sheldrake, 1973). Whereas import can occur by simple diffusion, export likely necessitates the activity of a transport protein (Kramer, 2006;Binenbaum et al., 2018). Of the more than 130 different GAs existing in nature (MacMillan, 2001) a handful has been tested in various *in vitro* transport assays and shown varying membrane permeation (Saito et al., 2015;David et al., 2016;Kanno et al., 2016;Tal et al., 2016). In particular, products of the C13-hydroxylation pathway appear less membrane permeable than their non C13-hydroxylated counterparts (Binenbaum et al., 2018). To investigate this in more detail, we exposed water injected control oocytes to a 50 µM equimolar mixture of eight different GAs (GA1, GA3, GA4, GA7, GA8, GA9, GA19 and GA24) at three different external pH values (pH 4.7, pH 5.3 and pH 6.0) for 60 min and subsequently quantified internalized GA by LC-MS/MS analysis of oocytes homogenates (Figure 1a). The eight GAs represent intermediate, bioactive and catabolic GA species and characteristically contain a varying number of oxygen atoms (Figure 1b). Four of the GAs (GA4, GA7, GA9 and GA24) appeared to permeate the membrane as they were detected in oocytes at all external pH values whereas the remaining four GAs (GA1, GA3, GA8 and GA19) were not detected in oocytes regardless of the external pH. Increasing the external pH lowered the accumulation of membrane permeable GAs. Based on the quantification, we categorized three of the permeating GAs (GA4, GA7 and GA24) as moderately permeating (oocyte GA concentration ≤ external concentration at pH 5.3 and 6.0, respectively) and one (GA9) as highly permeating (internal concentration 20-80 fold over external concentration at pH 6 to 4.7). From this dataset, we draw a simple correlation between number of oxygen atoms and membrane permeability (Figure 1c). Irrespective of oxygen positions on the GA backbone or the functional group in which they participate, GAs with 6 or more oxygens (GA1, GA3, GA8 and GA19), did not permeate the *Xenopus* oocyte membrane at any pH. In comparison, GAs with 5 oxygens (GA4, GA7 and GA24) permeate moderately and GAs with only 4 oxygens (GA9) permeate to a high degree. Thus, background accumulation due to diffusion of protonated membrane permeating GAs can - not surprisingly - be minimized by increasing external pH in transport assays. It should be emphasized here that we only investigated membrane permeability toward various GAs in *Xenopus* oocytes and that similar permeability/background uptake cannot necessarily be inferred for other heterologous hosts such as yeast. For example, GA3, which does not permeate oocytes appears to accumulate weakly in yeast (Saito et al., 2015;David et al., 2016;Kanno et al., 2016;Tal et al., 2016).

**Figure 1.**
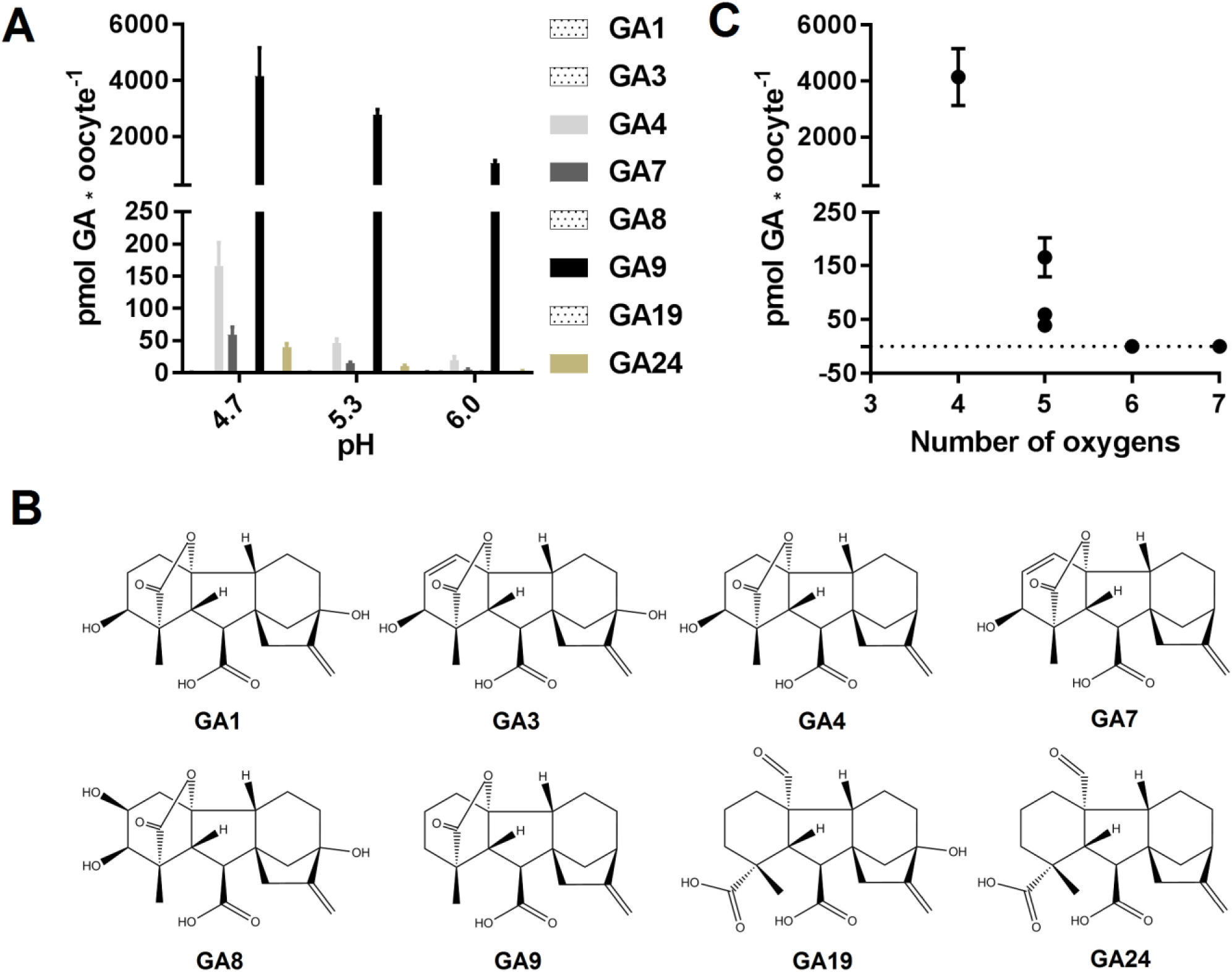
GA membrane permeability is a function of pH and oxygen content. (**A**). Mock expressing oocytes (n = 7-9) were exposed to a mix of 50 µM GA1, 50 µM GA3, 50 µM GA4, 50 µM GA7, 50 µM GA8, 50 µM GA9, 50 µM GA19, and 50 µM GA24 in pH 4.7, 5.3 and 6.0 for 60 min and GA content was quantified by LC-MS/MS. (**B**). Chemical structures of the tested GAs. (**C**). GA content of mock injected oocytes at pH 4.7 as a function of GA oxygen content.

### 3.2 Determining optimal pH for GA transport assays in *Xenopus* oocytes expressing NPF proteins

To determine a suitable pH for GA transport assays in *Xenopus* oocytes, we used NPF3.1 as a study case. This transporter imports a wide range of GA species when expressed in *Xenopus* oocytes and assayed at pH 5 (Tal et al., 2016). However, no difference to mock can be seen when exposed to highly permeating GAs such as GA9 or GA12. One explanation could be that the strong diffusion of GA9 into oocytes may mask any transport activity (Figure 1a). We have previously shown that transport activity of NPF3.1 is undetectable at pH 7 (Tal et al., 2016). Here, we test the activity of NPF3.1, in a 180 min time-course uptake assay towards 50 µM of the non-permeating GA3 at pH 5 and 6 (Figure 2a). At pH 5, NPF3.1-mediated accumulation of GA3 reached equilibrium after approximately 120 min incubation (∼1.5 fold x external medium concentrations). At pH 6, NPF3.1- mediated GA3 uptake also reached equilibrium after two hours incubation albeit the equilibrium level was approximately 75 % lower than at pH 5. For comparison, we tested the membrane permeability of GA9 in a 180 min time-course assay in water injected control oocytes (Figure 2b). Accordingly, seeking a compromise between reduced diffusion and reduced transport activity, we retested whether NPF3.1-mediated transport of GA9 into oocytes could be detected at pH 5.5 and pH 6. No difference could be seen to mock oocytes when assayed at pH 5.5 (i.e. both accumulated GA9 to equally high levels). In comparison, when assayed at pH 6, NPF3.1 expressing oocytes accumulate higher amounts of GA9 compared to water injected control oocytes (Figure 2c). Thus, for characterizing NPF protein mediated transport of highly permeating GAs in *Xenopus* oocytes, it can be advisable to adjust external media to pH values >5.5 to reduce diffusion to an extent where the contribution of transport activity is distinguishable. As the transport activity is dramatically reduced at pH 6 (Figure 2a) we opted for exploring NPF3.1 transport activity towards our selection of non-permeating GAs at pH 5, moderately permeating GAs at pH 5.5 and highly permeating GAs (including GA12) at pH 6. We detected significantly higher accumulation compared to water injected control oocytes for all non-permeating, moderately permeating and highly permeating GAs at pH 5, pH 5.5 and pH 6, respectively (Supplementary Figure 1). Thus, we established optimized conditions for quantitative GA transport assays capable of detecting NPF3.1-mediated accumulation of eight different GAs including the highly membrane permeating species.

**Figure 2.**
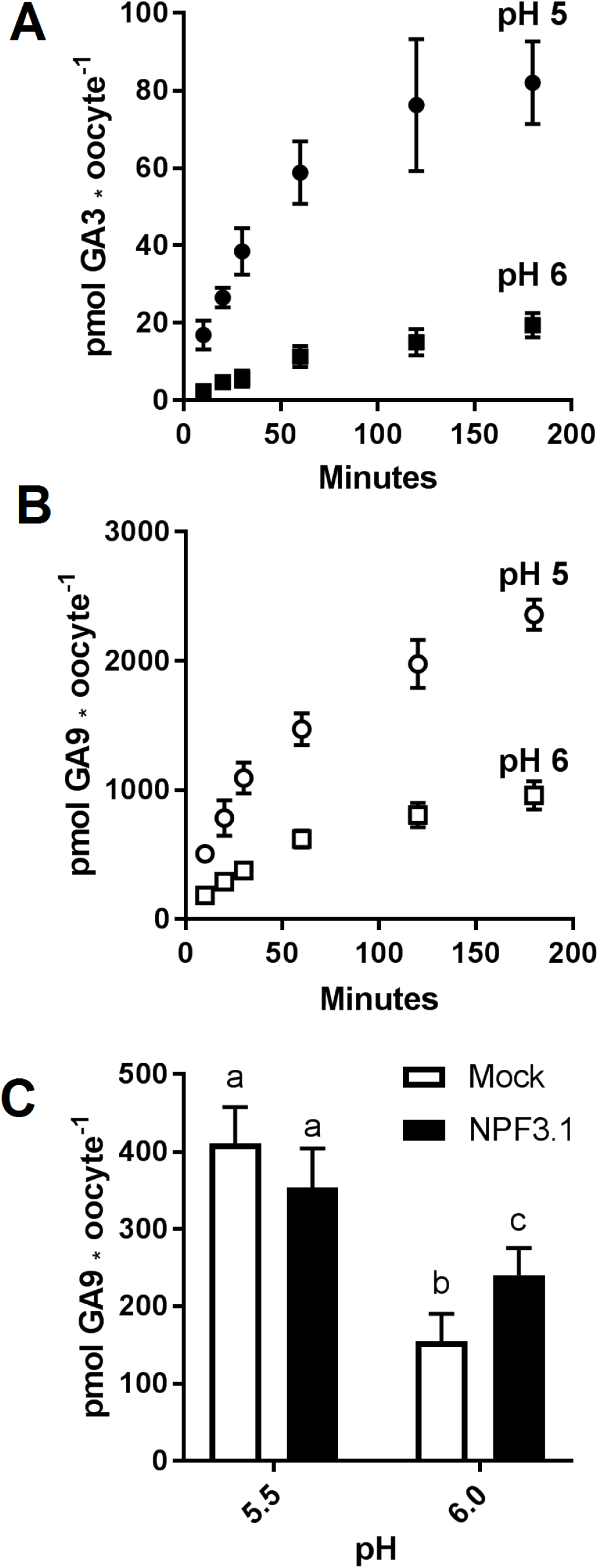
Assessment of GA permeability and NPF transporter function. (**A**). Time course of NPF3.1 mediate GA3 transport over 180 min. NPF3.1 expressing oocytes were exposed to 50 µM membrane non-permeable GA3 in kulori pH 5 or 6. At each time point, oocytes (n = 6) were taken out for analysis. (**B**). Time course of GA9 membrane penetration over 180 min. Mock expressing oocytes were exposed to 50 µM GA9 in kulori pH 5 or 6. At each time point, oocytes (n = 6) were taken out for analysis. (**C**). NPF3.1 mediated GA9 transport at pH 5.5 and 6. Oocytes (n = 6) were exposed to 50 µM GA9 in kulori (materials and methods) pH 5.5 or 6 for 60 min. Oocyte GA content was analyzed using LC-MS/MS. Statistical assessment was performed with Holm Sidak two-way ANOVA p = 0.01.

### 3.3 The proton potassium antiport activity portraits an efflux transporter artefact

In theory, passive diffusion of moderately permeating GA into oocytes allows screening for exporting transport proteins. For example, if an oocyte expressing a candidate exporter accumulates less GA4 compared to mock oocytes, it could indicate that a portion of the GA4 molecules that have diffused into the transporter expressing oocytes can move out of the oocyte through the transporter. In our preparatory phases we exposed parts of our NPF library to a mixture of phytohormones at external pH 5. The mixture included JA, which highly permeates oocyte membranes at acidic pH. Strikingly, three related transporters, NPF7.1-7.3 all accumulated 75-85% less JA compared to water injected control oocytes (Figure 3a). Our excitement about having identified potential JA exporters was, however, curbed by several factors. Firstly, in an attempt to determine the transporters’ substrate preference, we exposed NPF7.3 expressing oocytes to five different phytohormones with lipophilic weak acid properties (GA4, ABA, JA, JA-Ile and OPDA). Despite the diversity in chemical structures, NPF7.3 expressing oocytes accumulated significantly less of all five phytohormones compared to water injected control oocytes (Figure 3b). Thus, either this putative exporter appeared to possess the same enigmatic multi-substrate specificity towards phytohormones as described for other NPF proteins (Kanno et al., 2012;Chiba et al., 2015;Tal et al., 2016) or the reduced accumulation within oocytes was possibly due to indirect effects. In the context of *in vitro* transport assays, the accumulation equilibrium generated by the ion-trap mechanism is expectedly sensitive to small pH changes on either side of the membrane (Rubery and Sheldrake, 1973). Recently, NPF7.3 was shown to exhibit a non-electrogenic proton/potassium antiport activity when expressed in oocytes. However, the effect of the imported protons on internal oocyte pH was not investigated (Li et al., 2017). This prompted us to measure intracellular pH in oocytes expressing NPF7.3 using a proton-selective electrode in a three-electrode voltage clamp setup. Oocytes were clamped at a membrane potential of 0 mV and perfused with kulori at pH 7.4. This was followed by 400 secs perfusion with ekulori buffer pH 5.0 and then back to pH 7.4 for additional 400 secs, while the intracellular pH was monitored continuously. Compared to water injected control oocytes, NPF7.3 expressing oocytes displayed a significant 0.24 pH-units higher cytosolic pH in buffers with pH 7.4. However, within 400 secs intracellular pH markedly dropped by 0.5 pH units when external pH was lowered from 7.4 to 5.0, reaching a cytosolic pH 0.26 units lower than in water injected control oocytes (Figure 3c). During prolonged exposure (60 min) to ekulori buffer pH 5.0 intracellular pH even decreased to a stable pH of approximately 6.3, that completely reversed when re-subjected to pH 7.4 (Supplementary Figure 2). In comparison, oocytes individually expressing a selection of other NPF members; NPF2.5, NPF2.13, NPF3.1, NPF4.1, NPF4.3 and NPF4.6 (Figure 4) did not display a drop in intracellular pH that was significantly different than water injected control oocytes. Next we tested whether a decrease in intracellular pH of this magnitude affects the accumulation of membrane permeating phytohormones. Due to its rapid diffusion into oocytes, we used JA as a representative example of diffusing phytohormones. First, we investigated JA accumulation in water injected control oocytes wherein pH was lowered by concentrated MES/TRIS buffer (pH 5.7) injection resulting in an oocyte intracellular pH of ∼6.25 that was stable for approximately 30 min (Supplementary Figure 3). This procedure mimicked the pH lowering effect of NPF7.3 Next we exposed these oocytes to 100 µM JA for 20 min. In comparison, to oocytes that were not injected with the pH-lowering buffer, oocytes with the buffer-controlled intracellular pH of ∼6.25 accumulated significantly less JA (Figure 3d). In contrast, when we stabilized intracellular pH of NPF7.3 expressing oocytes to ∼7.5 via buffer injection, these oocytes accumulated JA to the same extent as in mock oocytes with no buffer injection (Figure 3d). These results strongly suggest that altered accumulation of phytohormones in NPF7.3 expressing oocytes is an indirect effect of the proton influx and concomitant lowering of intracellular pH.

**Figure 3.**
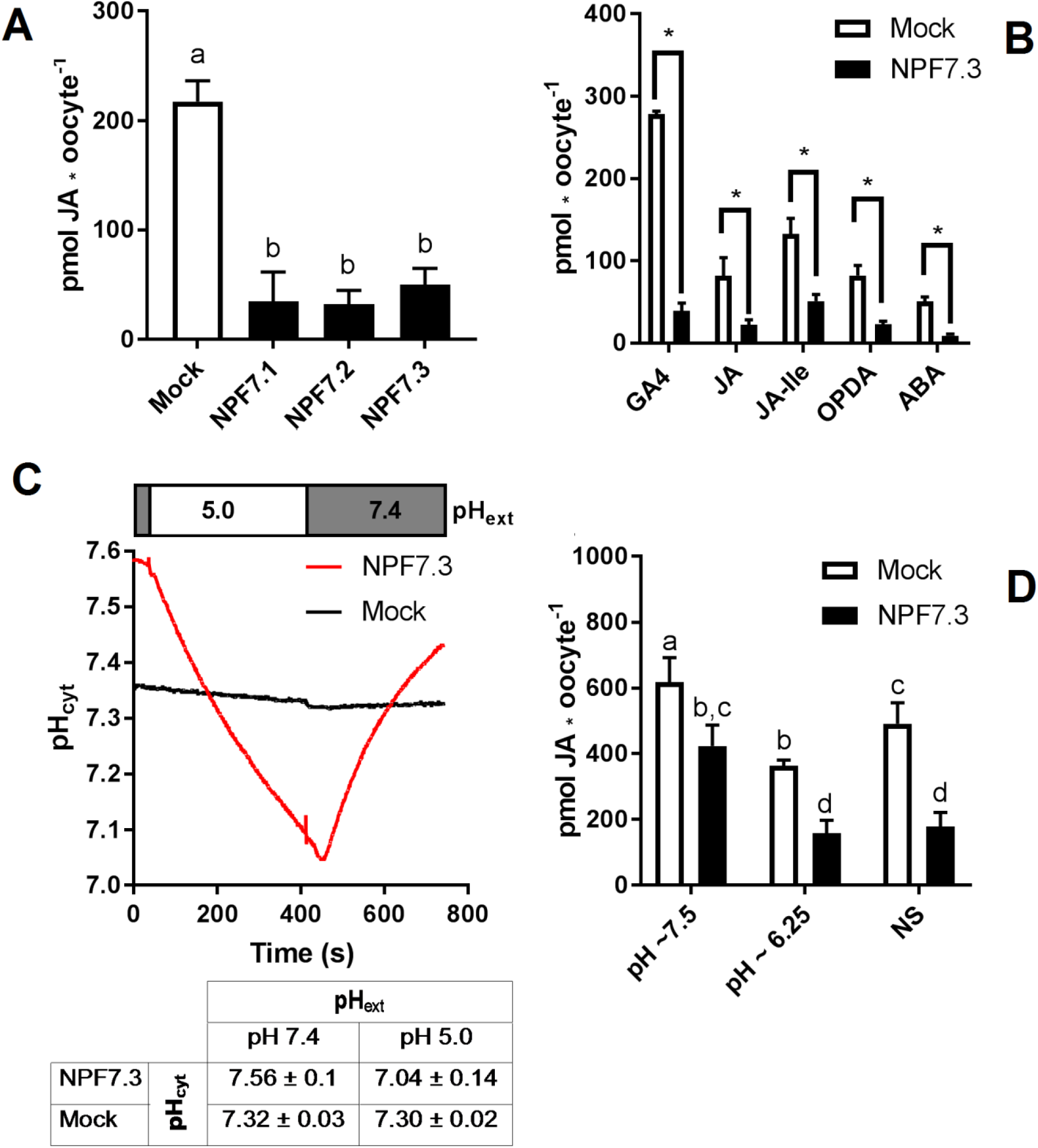
The proton potassium antiport function of NPF7.3 influences the ion-trap mechanism of membrane permeable weak acids. (**A**). Oocytes were exposed to 100 µM JA in kulori pH 5 for 60 min and analyzed in 3 technical replicates of 5 oocytes. Statistical assessment was performed with Holm Sidak one-way ANOVA p = 0.01. (**B**). Lower accumulation of membrane permeable phytohormones in NPF7.3 expressing oocytes compared to mock. Oocytes were exposed to 100 µM phytohormone in kulori pH 5 for 60 min (JA, JA-Ile, OPDA, ABA) or 90 min (GA4) and analyzed in 3-4 technical reps of 4-5 oocytes. Statistical assessment was performed with two-tailed t-tests p =0.05 (**C**). Upper panel: Internal oocyte pH measured using three-electrode voltage-clamp of mock vs NPF7.3 expressing oocytes. Starting in ekulori pH 7.4, the external buffer was changed to ekulori pH 5.0 for 400 secs followed by ekulori pH 7.4 for another 400 sec. pH was measured continuously at a membrane potential of 0 mV. Representative measurements are shown. Lower panel: Mean values of cytosolic pH in mock or NPF7.3 expressing oocytes in standard ekulori buffer pH 7.4 or 400 secs after incubation in ekulori pH 5.0 (n ≥ 3, mean ± SD). (**D**). JA content at defined cytosolic pH. Oocytes injected with 0.5 M TRIS 50 mM EGTA adjusted to pH 7.7 with 0.5 M MES (pH stabilized at ∼7.5), 0.5 M MES 50 mM EGTA adjusted to 5.7 with 0.5 M TRIS (pH stabilized at ∼6.25) or water (not stabilized: NS) was exposed to 100 µM JA in kulori pH 5 for 20 min and analyzed in 2 x 3 technical reps of 4-5 oocytes. Oocyte phytohormone content was analyzed using LC-MS/MS. Statistical assessment was performed with Holm Sidak two-way ANOVA p = 0.01.

**Figure 4.**
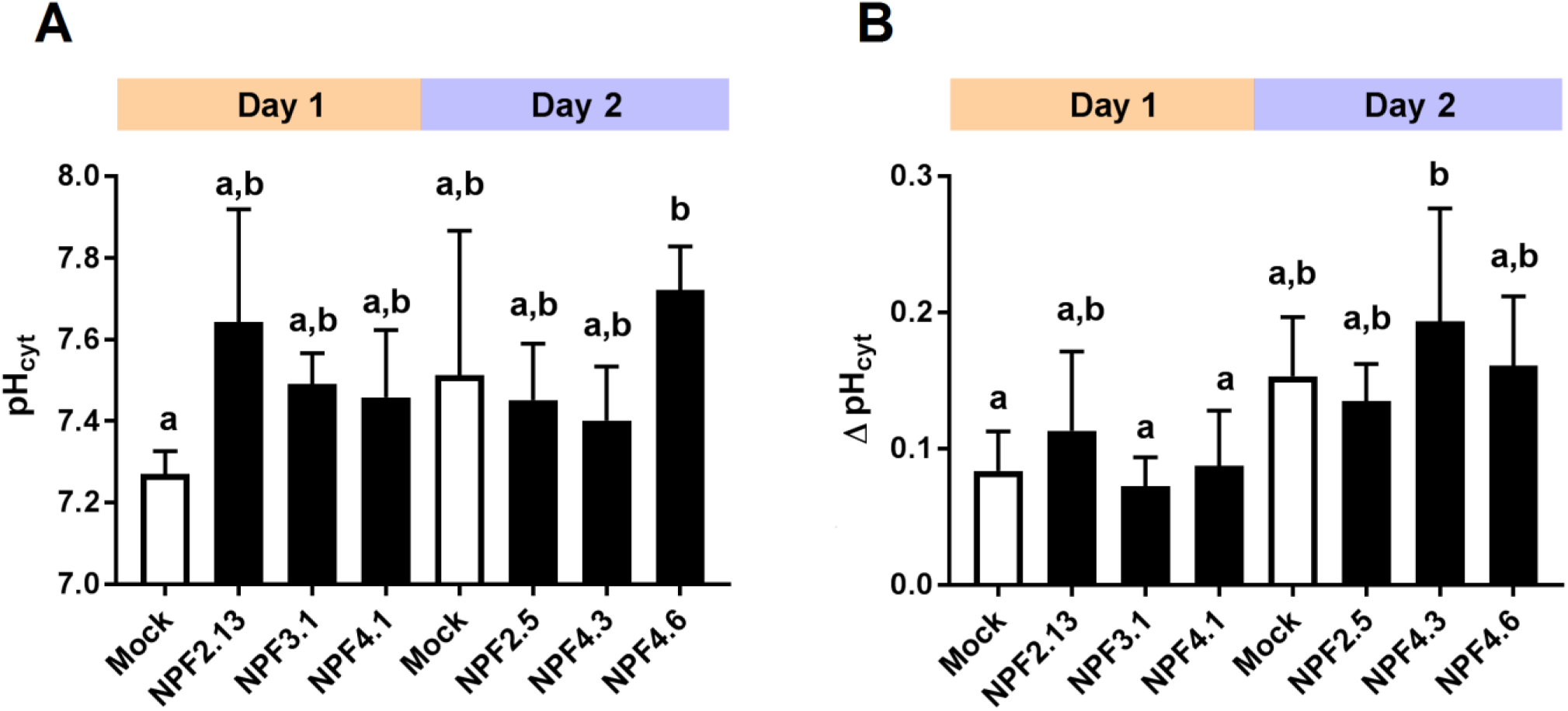
Transporter expression and oocyte maturity influence internal oocyte pH. Internal oocyte pH measured using three-electrode voltage-clamp of mock versus expressing oocytes. (**A**). after 30 min resting in ekulori pH 7.4 and additional 5 min clamped at 0 mV with pH 7.4 ekulori perfusion. (**B**). Change in internal pH after additional 5 min perfusion with pH 5 ekulori. Statistical assessment was performed with Holm Sidak two-way ANOVA p = 0.05.

### 3.4 A quantitative screen of the *Arabidopsis* NPF proteins

To ascertain the *in vitro* based functional annotation of NPF GA transport activity, we used the optimized GA transport conditions to quantitatively screen for GA transport activity towards non-permeating and moderately permeating GAs in oocytes. Therefore, cRNA for translation of all 53 *Arabidopsis* NPF members were injected individually into *Xenopus* oocytes and after three days of expression oocytes were exposed for 60 min at pH 5.5 to mixtures of bioactive GAs (50 µM GA1 and 100 µM GA4) (Yamaguchi, 2008), a product of catabolism (50 µM GA8) (Yamaguchi, 2008), a biosynthesis intermediate (50 µM GA19) (Yamaguchi, 2008), phloem transported (50 µM GA24) (Regnault et al., 2015) and the seed specific (50 µM GA3) (Derkx et al., 1994). The data is displayed in two figures (Figure 5 and Figure 6). Figure 5 displays transport activities for non-permeating GAs (GA1, GA3, GA8 and GA19), whereas Figure 6 displays transport activities of the moderately permeating GAs (GA4 and GA24). Un-normalized data is included in (Supplementary Figure 4). From subclade NPF4, only NPF4.1 and NPF4.6 transported non-permeating GAs, where NPF4.6 accumulated only ∼15% of GA compared to NPF4.1 levels. The only other transporters displaying uptake activities of similar magnitude as NPF4.1 were NPF2.5 and NPF3.1. NPF3.1 imported approximately 40 % of NPF4.1 levels. In comparison, NPF1.1, NPF2.3, NPF2.4, NPF2.7, NPF2.12 and NPF2.13 imported GAs to approximately 10-15% of NPF4.1 levels. Thus, no significant uptake was detected in oocytes expressing any NPF member from subclades 5-8, indicating that GA transporters cluster within subclades 1-4.

**Figure 5.**
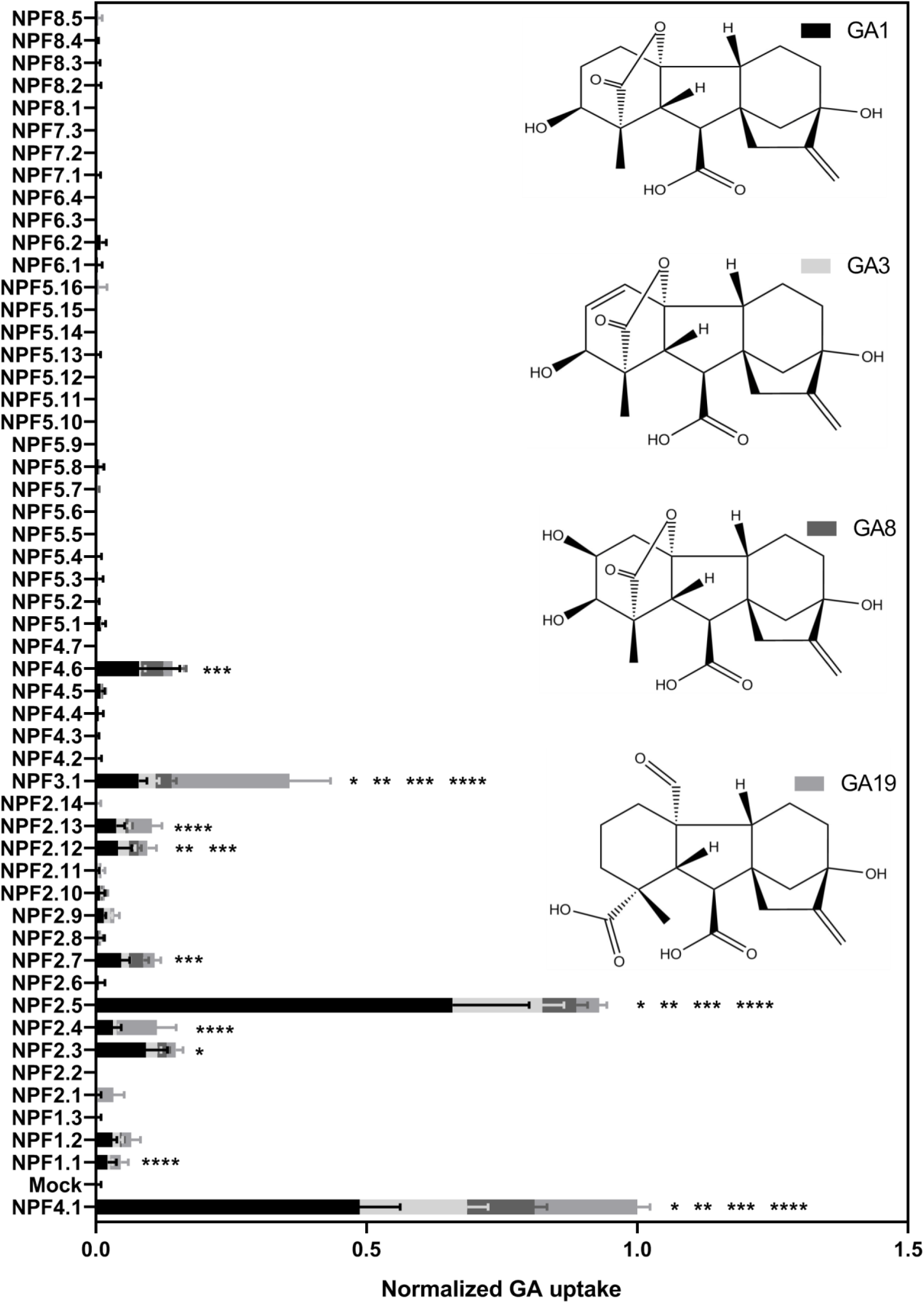
Quantitative screen of the NPFs for GA transport of not membrane non-permeable GAs. Due to logistic considerations, the 53 NPF members were screened in two portions on the same day and normalized to the transport of NPF4.1 of membrane non-permeable GAs. NPF4.1 was chosen for normalization as it is a well characterized GA transporter (Kanno et al., 2012;Saito et al., 2015;Tal et al., 2016). For each transporter the proportion of each GA imported into oocytes is given in proportion to total imported amount of GAs. Statistical significant transport (Holm Sidak one-way ANOVA p = 0.05) is indicated with one asterisk for GA1, two asterisks for GA3, three asterisks for GA8 and four asterisks for GA19.Two assays of 28 and 29 genes, respectively, were performed on same day on the same oocyte batch. Both assays included NPF3.1, NPF4.1 and Mock in order to normalize. Oocytes (n = 5-6) were exposed to a mix of 50 µM GA1, 50 µM GA3, 100 µM GA4, 50 µM GA8, 50 µM GA19 and 50 µM GA24 in pH kulori 5.5 (Materials and Methods) for 60 min. Oocyte GA content was analyzed using LC-MS/MS.

**Figure 6.**
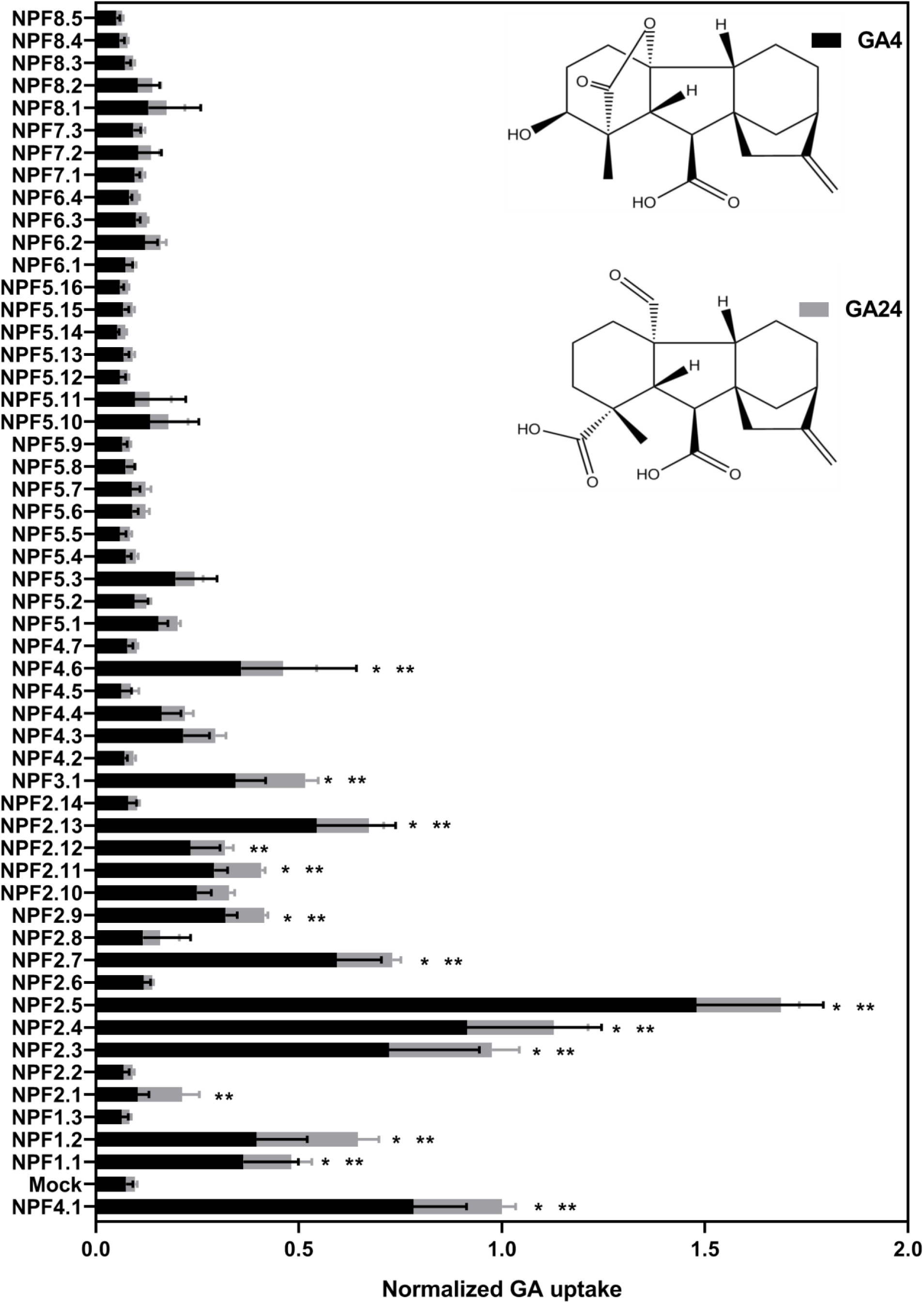
Quantitative screen of the NPF proteins for GA transport of membrane permeable GAs. Due to logistic considerations, the 53 NPF members were screened in two portions on the same day and normalized to the transport of NPF4.1 of membrane permeable GAs. NPF4.1 was chosen for normalization as it is a well characterized GA transporter (Kanno et al., 2012;Saito et al., 2015;Tal et al., 2016). For each transporter the proportion of each GA imported into oocytes is given in proportion to total imported amount of GAs. Statistical significant transport (Holm Sidak one-way ANOVA p = 0.05) is indicated with one asterisk for GA4 and two asterisks for GA24. Two assays of 28 and 29 genes, respectively, were performed on same day on the same oocyte batch. Both assays included NPF3.1, NPF4.1 and Mock in order to normalize. Oocytes (n = 5-6) were exposed to a mix of 50 µM GA1, 50 µM GA3, 100 µM GA4, 50 µM GA8, 50 µM GA19 and 50 µM GA24 in pH kulori 5.5 (Materials and Methods) for 60 min. Oocyte GA content was analyzed using LC-MS/MS.

Interestingly, our screen indicates different substrate preferences among the identified GA transporters. However, due to a necessity for more rigorous characterization we did not score differences in preference statistically. However, some clear potential preferences can be seen for a few genes. For example, NPF3.1 appears to prefer the biosynthesis intermediate GA19, whereas NPF2.5 shows a strong preference for the bioactive GA1.

Transport data on the moderately permeating GAs identifies significant transport activity for more putative GA transporters than uptake on non-permeating GAs. Significant transport is still confined to subclades 1-4 but with more members of subclades 1 and 2 identified as potential GA transporters, namely (NPF1.2, NPF2.1, NPF2.9, and NPF2.11).

### 3.5 Delineating potential substrate binding residues

The uptake results presented above (Figure 5 and Figure 6) shows that the apparent GA transport activities are confined to a subset of NPF proteins from subclades 1-4. If we assume that GA-transporting NPF proteins must have residues that confer GA selectivity compared to non-transporting NPF proteins; a pertinent question is whether the identified NPF members share these residues to form identifiable GA specific motifs.

The canonical MFS transporter structure consists of two six-helix bundles connected by a long cytosolic loop; the interface between these two bundles contains the residues that interact with the transported substrate and thus define the substrate specificity. The crystal structure of NPF6.3 is an excellent example of an MFS structure (Parker and Newstead, 2014;Sun et al., 2014). During its transport cycle the transporter will alternate between conformations with the substrate-binding site open to the extracellular side, be occluded, or open to the cytoplasm (Jardetzky, 1966;Abramson et al., 2003;Huang et al., 2003). POT family members and NPF members are closely related (Fei et al., 1994;Léran et al., 2014;Jørgensen et al., 2015). Several POT crystal structures have been co-crystallized with a number of different peptides or peptidomimetic drugs (Doki et al., 2013;Guettou et al., 2014;Lyons et al., 2014;Newstead, 2015;2017). All peptides bind at relatively equivalent positions between the two six-helix bundles. To identify potential specificity determining residues of the NPF proteins we utilized POT structures crystallized in complex with the peptidomimetic drug alafosfalin (PDB: 4IKZ and 4LEP) (Doki et al., 2013;Guettou et al., 2013). Surface exposed residues within an 8 Å sphere around alafosfalin were selected; additionally, residues predicted to be surface exposed in an outward-facing conformation, as judged from comparisons with the FucP and YajR structures (PDB: 3O7Q and 3WDO) (Dang et al., 2010;Jiang et al., 2013) were also selected; yielding a total of 51 residues (Figure 7, Supplementary Video 1. From this set of residues, we created sequence logos for the non-membrane permeating GA (GA1, GA3, GA7, GA19) transporting-versus non-GA transporting *Arabidopsis* NPF proteins (GA(+) and GA(-), respectively in Figure 7) in order to highlight any preferred positions (Crooks et al., 2004). Only a few distinct positions were observed; hereunder arginine in position 16, hereafter denoted Arg^Pos16^ (Lys164 of NPF6.3), Ser^Pos28^ (Trp353 of NPF6.3) and Gln^Pos31^ (Leu359 of NPF6.3) (Figure 7).

**Figure 7.**
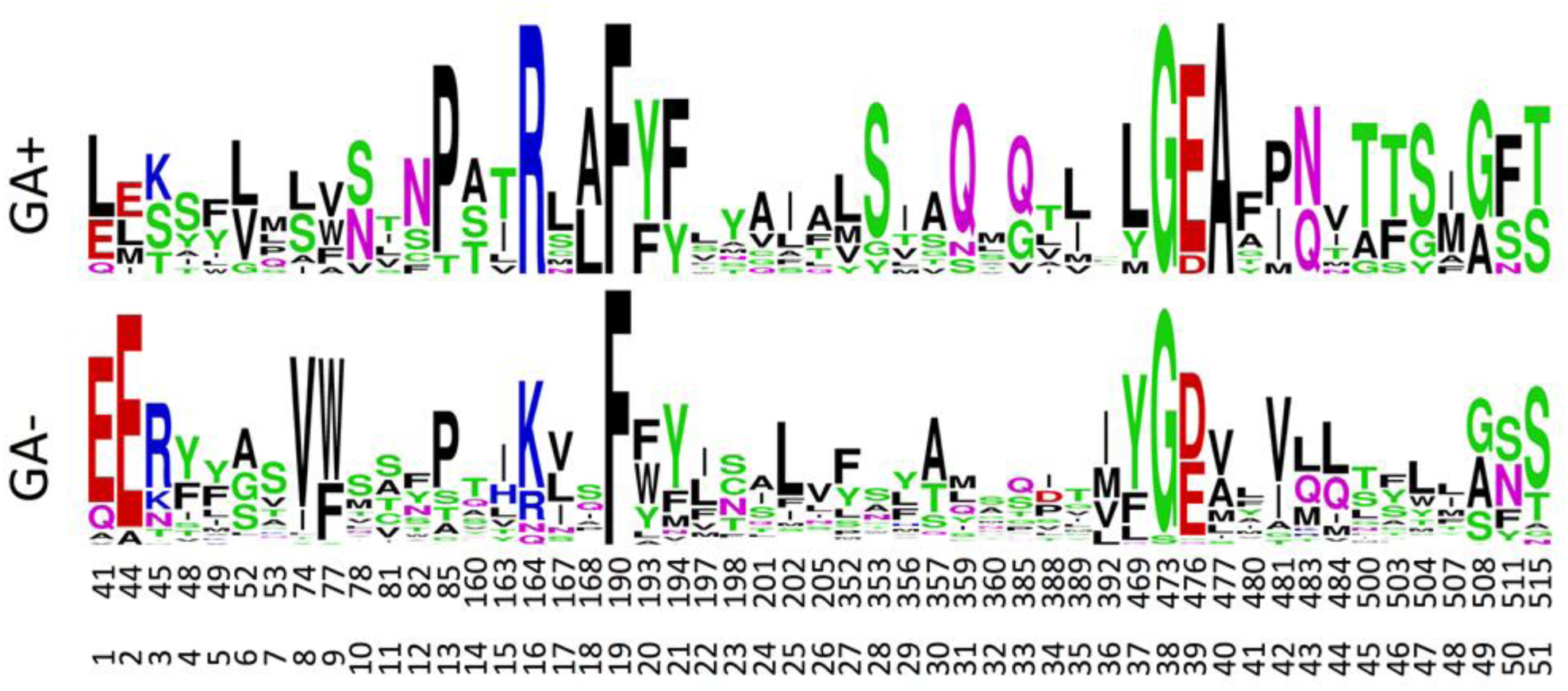
Sequence logos for the GA transporting (GA+) versus GA non-transporting (GA-) NPF proteins. Numbers correspond to the amino acid position of *Arabidopsis* thaliana NPF6.3 and the relative position of the binding site residues. The figure was made in the WebLogo program (Crooks et al., 2004).

For assessing the physiochemical environment in the substrate binding cavity in NPF transporters the same set of residues were converted to the position-dependent numerical descriptors, z-scales, developed by (Hellberg et al., 1987). The original z-scales descriptors have been derived by a PCA of 29 physicochemical variables describing the properties of the 20 natural amino acids, and represent the hydrophilicity (*z*_*1*_), steric properties (*z*_*2*_) and polarity of the amino acid (*z*_*3*_). The z-scales have successfully been used in several proteochemometric studies, for instance to model HIV protease resistance or to alignment-independently classify G-coupled receptors and more (Lapinsh et al., 2002;van Westen et al., 2013;Cortés-Ciriano et al., 2015). The 51 sequences times 51 residues matrix was converted to a 51 sequences times 153 z-scales matrix, which was subjected to a PCA.

The PCA only shows four major clusters indicating shared physicochemical properties in the substrate binding cavity in genes across phylogenetic subclades. Three of the clusters are clearly defined whereas the fourth is larger and more diffuse. Cluster I, contains six NPF2 subclade proteins that include four GA(+) transporters and two GA(-) transporters. Cluster II contains eleven NPF proteins, four GA(+) and seven GA(-) transporters, from the NPF1, NPF2 and NPF3 subclades. Cluster III, contains solely GA(-) transporters from the NPF5 subclade. Finally, cluster IV contains the remaining NPF proteins including the two GA(+) transporters, NPF4.1 and NPF4.6 (Figure 8). Glucosinolate transporting NPF proteins (NPF2.9-2.13 and NPF3.1) (Nour-Eldin et al., 2012;Jørgensen et al., 2017) and peptide transporting NPF proteins (NPF5.2 and NPF8.1-NPF8.3) (Supplementary Figure 5) (Frommer et al., 1994;Dietrich et al., 2004;Hammes et al., 2010) group in two tight groups in distinct clusters in the PCA plot (Figure 8), and both clusters contain additional genes (cluster II and IV, respectively). Similar to GA(+) transporters, nitrate transporters are scattered in more clusters (Supplementary Figure 6).

**Figure 8.**
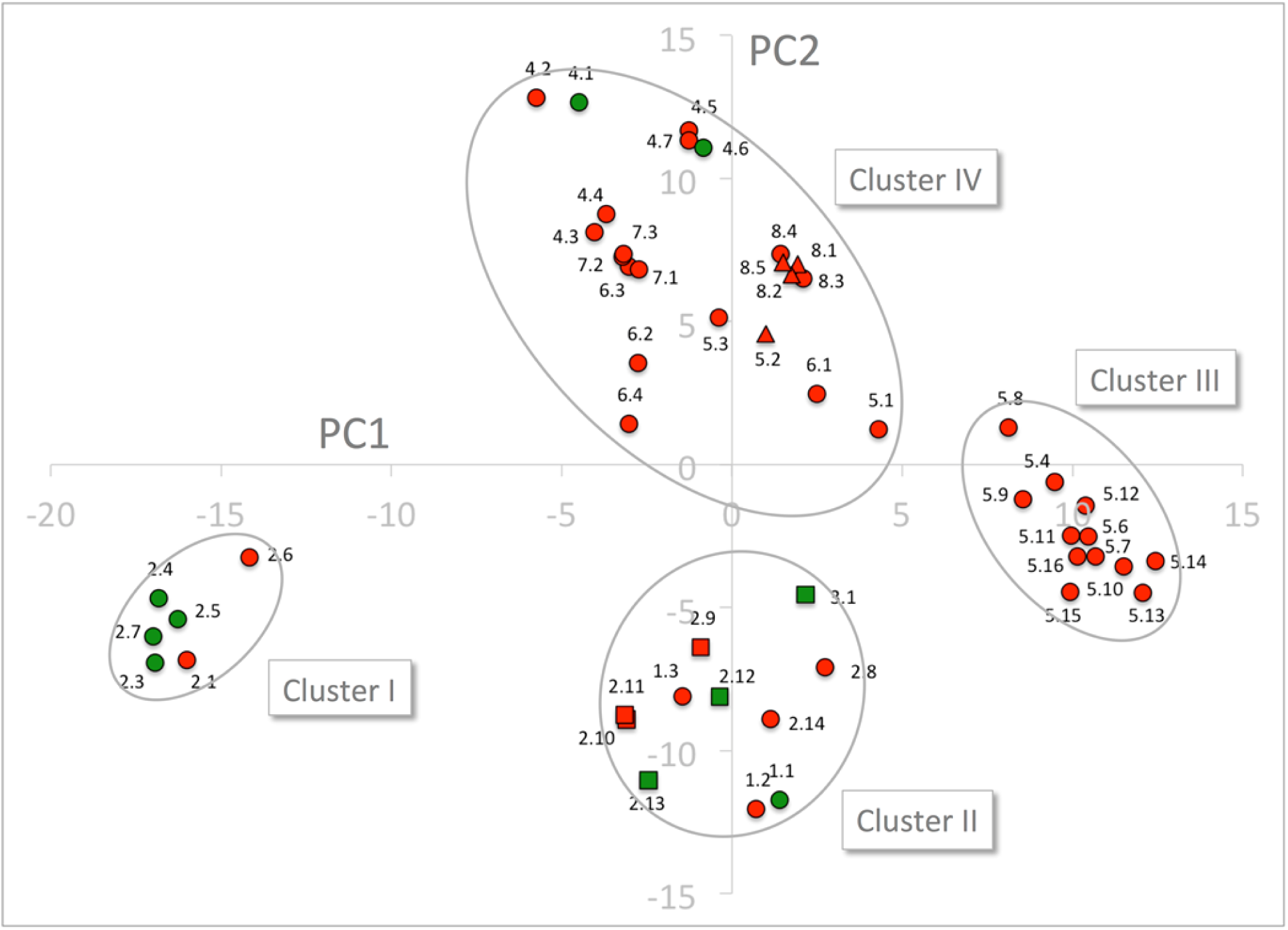
PCA of the 51 NPF sequences expressed by z-scales. GA transporting transporters are shown as green dots and squares and GA non-transporting transporters as red dots, triangles and squares. Glucosinolate transporting genes and peptide transporting genes are shown as squares and triangles, respectively. The four clusters are marked by ellipses. PC1 and PC2 refer to the first and second principal components, respectively.

Cluster I is composed exclusively of genes belonging to the NAXT subclade whose members all lack the ExxE[K/R] motif, i.e. the negatively charged Glu^Pos1^ (Residue 41 of NPF6.3), Glu^Pos2^ (Residue 44 of NPF6.3) and the positively charged Arg/Lys^Pos3^ (Residue 45 of NPF6.3). This motif is otherwise conserved in the rest of the family except in the NPF7 clade (Segonzac et al., 2007;Jørgensen et al., 2015). To exclude bias by the presence/absence of the ExxE[K/R] motif, the entire PCA was repeated for only 48 residues (i.e. omitting positions 1-3). The 48 residue PCA yielded four clusters containing the same genes and a similar clustering as the PCA on the original 51 residues (Supplementary Figure 7). Thus, cluster I is not defined based on the absence of the ExxE[K/R] motif.

In order to identify characteristic properties for each of the four clusters, we created sequence logos for the individual clusters (Figure 9). Besides uniquely lacking the ExxE[K/R] motif, cluster I is the only cluster that includes Ala^Pos18^ (Ser168 of NPF6.3), Asn^Pos43^ (Gln483 of NPF6.3), Thr^Pos46^ (Leu503 of NPF6.3), Ser^Pos47^ (Leu504 of NPF6.3) and Phe^Pos50^ (Phe511 of NPF6.3) (Figure 9, red arrows; Figure 10, green residues). In comparison, only one residue is unique to cluster II, namely Asn^Pos12^ (Residue 82 of NPF6.3) (Figure 9, blue arrow; Figure 10, blue residue). It is interesting to compare the differences between cluster I/II versus cluster III/IV, since the majority of the GA transporters are found in cluster I and II. The sequence logos indicate residues that are uniquely shared between cluster I and II. For example, position 16 (Residue 164 in NPF6.3), is conserved as an Arg in cluster I and II and Lys in cluster III and IV. In addition, Gln^Pos31^ (Residue 359 of NPF6.3), Leu^Pos37^ (Residue 469 of NPF6.3) and Ala^Pos40^ (Residue 477 of NPF6.3) are characteristic for cluster I/II compared to cluster III/IV (Figure 9, black arrows; Figure 10, yellow residues).

**Figure 9.**
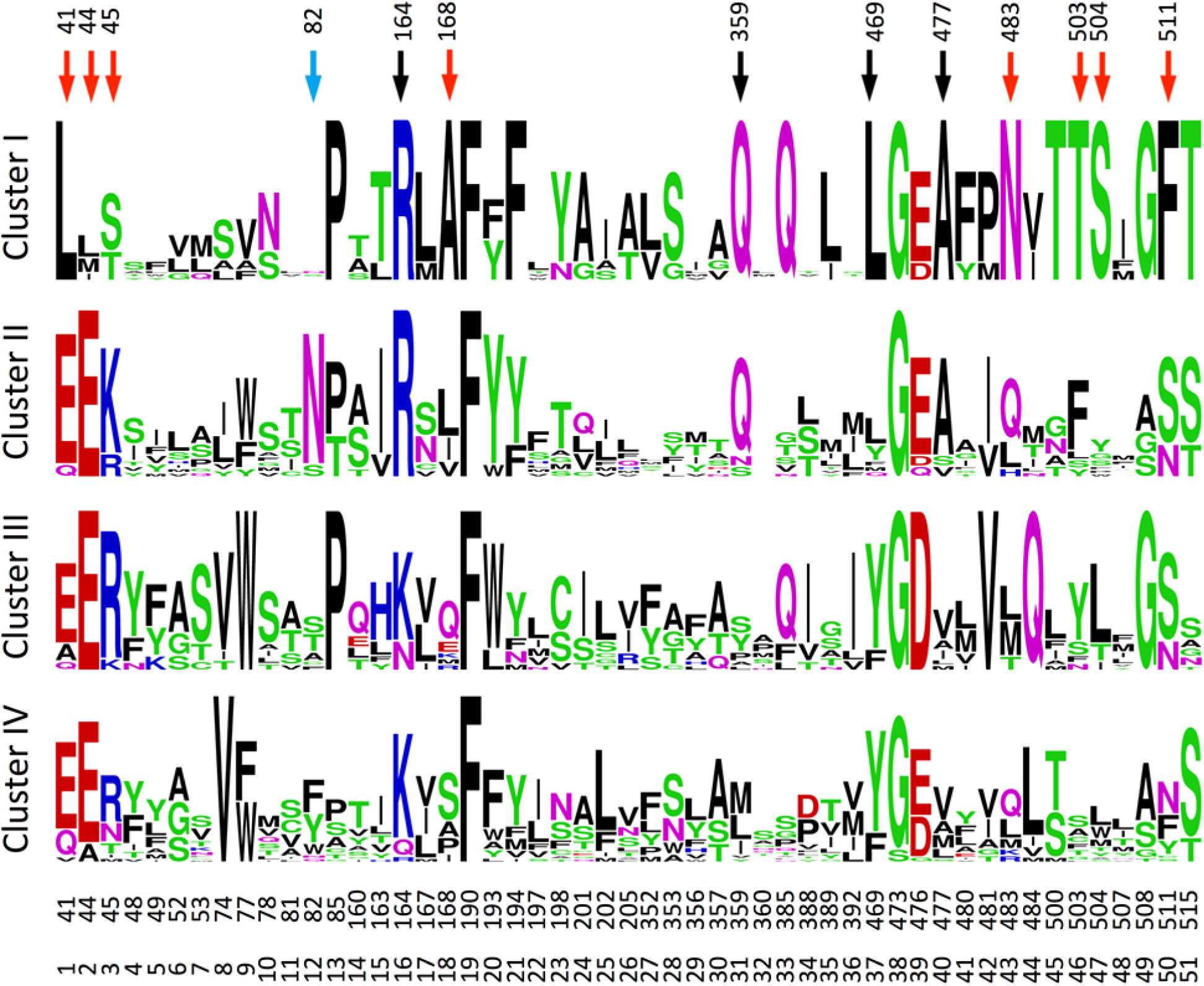
Sequence logos for the transporters in the four clusters. The figure was made in the WebLogo program (Crooks et al., 2004). Red, blue and black arrows mark positions unique for cluster I, cluster II and for cluster I and II, respectively.

**Figure 10.**
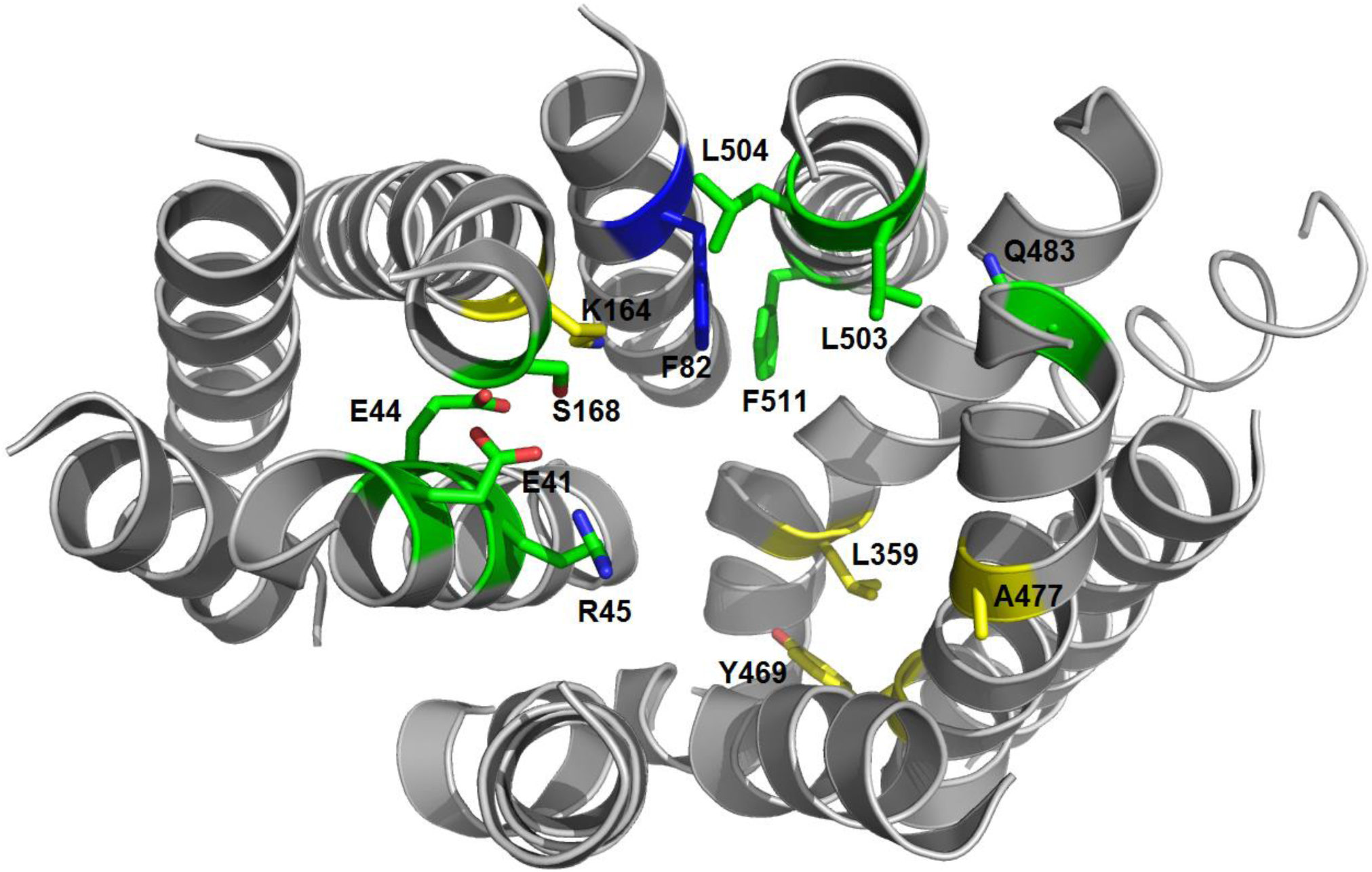
3D structure of the NPF6.3 (PDB: 4OH3 (Sun et al., 2014)). Residues unique for cluster I are shown as green sticks, the residue unique for cluster II is shown as a blue stick and residues unique for cluster I and II are shown as yellow sticks.

## 4 Discussion

Perhaps one of the most intriguing features of the NPF phytohormone transporters is the apparent multi-specificity towards phytohormones and other metabolites with distinct chemical structure (Corratge-Faillie and Lacombe, 2017;Wang et al., 2018). This phenomenon has previously prompted speculations in the NPF, providing a basis for the integration of environmental and physiological information linked to the relative availability of the different nutrients (Corratge-Faillie and Lacombe, 2017). However, to delve into this interesting notion of multi-substrate specificity it becomes pertinent to discuss how to distinguish between “real” and “non-” substrates for NPF transporters. In this context, our data provide some novel points for discussion.

Firstly, there appears to be indirect ways that NPF transporters can alter the distribution pattern of membrane permeating substrates with acid/base properties across membranes. For example, our characterization of NPF7.3 provides an alternative explanation to what we first perceived as multi-specificity towards different phytohormones. As NPF7.3 causes a profound reduction in internal oocyte pH when external pH is reduced from 7.4 to 5 (Figure 3c), the strength of the ion-trap mechanism is altered, and this affects the diffusion-based accumulation pattern of lipophilic phytohormones across the membrane regardless of their chemical structure. In this context, it is concerning that current *in vitro* data points to multi-substrate specificity for several NPF phytohormone importers (Supplementary Table 1). It remains to be investigated whether expression of NPF members in heterologous systems can cause increased internal pH that would lead to an increased ion-trap strength and thereby indirect uptake of many structurally unrelated membrane-permeating phytohormones. This could, for example, happen by the outward movement of protons or potentially by transport of buffer molecules through the transporters. Interestingly, *Streptococcus thermophilus* PepT1, a POT homolog of NPF transporters was recently co-crystallized together with the buffering reagent HEPES – binding within the cavity (Martinez Molledo et al., 2018). The efflux of protons has not yet been reported for any NPF protein nor has transport of HEPES to the best of our knowledge been investigated. We measured pH in oocytes expressing six different NPF proteins with GA import activity and could not detect altered pH in any of them (Figure 4). However, we cannot exclude that discrete and transient pH alterations may have eluded detection. In conclusion, when considering the results of transport assays involving membrane permeating lipophilic compounds with weak acid/base properties (such as most phytohormones), it is important to consider and exclude indirect factors that may cause changed distribution across the membrane. For example, if using *Xenopus* oocytes it is possible to inject a buffer with enough buffer capacity to maintain internal pH at neutral (Figure 3d).

Secondly, NPF2.10 and NPF2.11 (GTR1 and GTR2) transport a wide range of glucosinolates that all share the common glucosinolate core structure but with varying amino acid side chains (Nour-Eldin et al., 2012;Andersen et al., 2013;Jørgensen et al., 2017). Similarly, hPepT1 and hPepT2, two POT transporters involved in dietary peptide uptake in intestines, transport an immense range of peptidomimetics, indicating that a substrate binding cavity accommodating large variations over a common chemical core structure may be a common feature in the NPF (Biegel et al., 2006;Brandsch et al., 2008). Given this apparent plasticity in the substrate-binding cavity, it is conceivable that some NPF members (such as NPF2.10) may accommodate adventitious transport of a wide selection of metabolites (such as GAs) at low levels in heterologous systems. Discerning whether such transport activities represent “real” substrates ultimately, requires deeper insights into the relationship between structure and function of the NPF.

Here, we charted the GA substrate preference map of the *Arabidopsis* NPF proteins to elucidate whether it links to distinct structural features within the substrate-binding cavity. Such insights would represent first steps towards understanding the molecular basis of the selectivity of NPF transporters towards organic molecules and lead to an improved ability to predict substrate preference based on sequence information and 3D structure.

To date, almost all NPF-GA transporters have been identified using a qualitative and very sensitive substrate dependent growth-based screen in yeast using a two-component GA receptor-based yeast-two-hybrid system (Chiba et al., 2015). In addition, an *in vivo* approach monitored altered accumulation patterns of a fed bioactive fluorescein-conjugated GA3 in *Arabidopsis* transporter mutants. This approach identified NPF3.1 as a *bona fide* GA transporter with a role in accumulating GA in the endodermis (Shani et al., 2013;Tal et al., 2016). Despite the elegance of these approaches, their qualitative nature offers little distinction between transporters with varying transport activity and are, in their current versions, unable to detect transport of non-bioactive GA shown to undergo long distance transport *in planta* (Regnault et al., 2015;Binenbaum et al., 2018). Given its importance for downstream physiological and structure-function investigations, we found a need for critically re-assessing the *in vitro* data for GA substrate specificity among NPF transporters.

As a first step, we developed an optimized approach to screen for GA transporters among the *Arabidopsis* NPF members. Unlike previous approaches, the one presented here is quantitative, short-term, not limited to bioactive GAs, and gives preliminary insights into GA substrate specificity (Chiba et al., 2015). This is relevant as only some GAs species have been suggested to be transported long-distances in plants (Regnault et al., 2015). Additionally, compartmentalization has been suggested to play a regulatory role in GA signaling, thus transporters specific for either anabolic or catabolic GA species may be of interest (and can be identified in our screen) (Olszewski et al., 2002). In principle, by screening NPF transporters using a mixture of GAs that include moderately permeating GAs allowed us to screen for both import (increased accumulation) and export (reduced accumulation of permeating GAs) activity. However due to the potential ambiguity associated with interpreting accumulation of diffusing GA (described above), we focus on reporting and analyzing import activity only of non-diffusing GAs. In summary, our screen identified ten *bona fide Arabidopsis* NPF GA transporters belonging to subclades 1-4 that transported non-diffusing GAs in oocytes (Figure 5). In parallel, it is equally important to consider which genes our screen did not identify as GA transporters. In contrast to previous *in vitro* data, no members from subclades 5-8 were identified in our screen (Chiba et al., 2015). Notably, NPF1.3 appears to lack the large loop between TM6 and TM7 whereas NPF2.2 lacks a large region equal to parts of TM10 and TM11 of NPF6.3, thus it is likely that these two transporters are non-functional (data not shown). In support of the remaining negative uptake data not being false negatives e.g. due to lack of expression we note that 15 of these 39 transporters have previously been expressed functionally in *Xenopus* oocytes; NPF2.8, NPF2.10, NPF2.14, NPF5.5, NPF5.10, NPF5.11, NPF5.12, NPF5.16, NPF6.2, NPF6.3, NPF7.2, NPF7.3 NPF8.1, NPF8.2 and NPF8.3 (Tsay et al., 1993;Chiang et al., 2004;Chiu et al., 2004;Dietrich et al., 2004;Komarova et al., 2008;Li et al., 2010;Nour-Eldin et al., 2012;Léran et al., 2015;He et al., 2017;Jørgensen et al., 2017;Li et al., 2017). In conclusion, the quantitative short-term uptake screen presented here significantly reduces the number of potential *Arabidopsis* GA transporters from 25 to 10 and confines them to NPF subclades 1-4.

As a second step, we investigated whether molecular determinants of GA substrate specificity could be found among the transporter cavity exposed residues around the substrate binding sites. To identify these residues, we generated a map of all cavity exposed amino acids within an 8 Å sphere to the substrate binding site in all three conformations (outward open, occluded, inward open). This identified 51 positions in the plant NPF structures that we suggest as a general foundation for structure-function elucidation.

PCA analysis of physicochemical properties of the 51 residue subset and corresponding sequence logos did not reveal a clear set of features in the substrate-binding cavity that associated directly with all GA transporting NPFs. However, from our analyses a few residues did emerge that associated with a large portion of GA transporters. Namely, Arg^Pos16^, Gln^Pos31,^ Leu^Pos37^ and Ala^Pos40^ are either conserved differently in cluster I and cluster II in the PCA analysis of physicochemical properties compared to clusters III and IV or appear as distinct features in the family wide sequence logo analysis. As cluster I and II contains the majority of GA transporters it is possible that some of these residues are important for GA recognition. Incidentally, an investigation of YePepT from *Yersinia enterocolitica* identified Lys314 as a determinant of specificity towards negatively charged dipeptides (Boggavarapu et al., 2015). Lys314 of YePepT corresponds to position 31 (Leu359 of NPF6.3) in our alignment, which is highly and uniquely conserved as Gln^Pos31^ among the GA transporting NPF members of cluster I and II. Lastly, position 40 (Ala477 of NPF6.3), conserved as an Ala in cluster I and II, is located adjacent to a Glu which is highly conserved in all POTs where it plays a role in binding the N-terminus of peptides (Meredith et al., 2000;Jensen et al., 2012;Solcan et al., 2012). However, in summary as GA transporters NPF4.1 and NPF4.6 are located in cluster IV and that other specificities are known for members of clusters I and II, our analyses indicate that GA may be recognized differently by different NPF GA transporters.

It is intriguing that many of the *Arabidopsis* NPF proteins are able to transport nitrate, without any apparent binding site features substantiating these observations; nitrate is a small molecule and to some extent it might be comprehensible that it would be able to trigger the transport mechanism. Our assay results support previous observations and show that GA, a molecule much larger than nitrate, is essentially able to behave in the same way as nitrate and trigger transport mechanism without having any specific conserved binding site (Figure 8, Supplementary Figure 6). Human PepT1 and related bacterial transporters have been found to be highly promiscuous, nonetheless most if not all of these substrates contain structural remnants of peptides such as an N-or C-terminus, or a peptide bond. These groups presumably help anchoring the molecule in the substrate binding site for long enough to trigger transport. To our knowledge most NPF substrates known to date, as summarized in (Corratge-Faillie and Lacombe, 2017;Wang et al., 2018), contain a chemical group that can exhibit a negative charge but not a positive charge; it is conceivable that this negative charge anchors these substrates to a positively charged group in the binding site. Position 16 contains a conserved positive charge (Figure 7 and Figure 9) and has been shown to interact with the C-terminus of peptides (Jensen et al., 2014). Perhaps the negative charge of NPF substrates, including that of GAs, is what evolution has kept from the chemical properties of the peptides to ensure coupling the electrochemical proton gradient.

Lastly, it should be emphasized that functional annotations are of course very sensitive to false positive results. Despite our considerations of technical artifacts, it is still possible that our list of positive GA transporters is over-estimated. As exemplified by the GTRs which transport nitrate poorly and glucosinolates with high affinity (Wang and Tsay, 2011;Nour-Eldin et al., 2012;Jørgensen et al., 2017), we cannot exclude that a better substrate awaits to be identified for a number of the transporters in our list.

## 5 Abbreviations

ABA: Abscisic acid
GA: Gibberellic acid
JA: Jasmonic acid
JA-Ile: Jasmonoyl-isoleucine
MFS: Major Facilitator Superfamily
NPF: Nitrate transporter 1/Peptide transporter family
OPDA: Oxo-phytodienoic acid
PCA: Principal Component Analysis
POT: Proton-coupled Oligopeptide Transporter

## 6 Author contribution

NW optimized GA assays, performed the screen, participated in pH measurements, analyzed LC-MS/MS data contributed to the study design and wrote the paper based on a draft. MJ did the initial GA experiments including 4-Methylthio-3-butenyl and glycylsarcosine screen and contributed to pH measurements in *Xenopus* oocytes, NPF7.3 characterization, data analysis and the study design. HE made the multiple sequence alignment and defined the binding site residues. SL characterized NPF7.3 mediated phytohormone transport supported by ZB. TM participated in cytosolic pH measurements in *Xenopus* oocytes and data analysis. CC performed the LC-MS/MS analysis. MM synthesized the JA-Ile. DG participated in defining the NPF7.3 effect on the ion-trap mechanism. FJ performed PCA and clustering analysis. OM and HN advised and contributed to the study design. HN wrote the paper based on a draft. All authors discussed the results and commented on the manuscript.

## 8 Acknowledgements

We thank Patrick Achard for providing GA12 and Louise Svenningsen for technical assistance in the laboratory. This manuscript has been released as a Pre-Print at bioRxiv.

