## Supplementary figures and tables for "An Optimized Screen Reduces the Number of GA Transporters and provides Insights into NPF Substrate Determinants"

### *Supplementary Material*

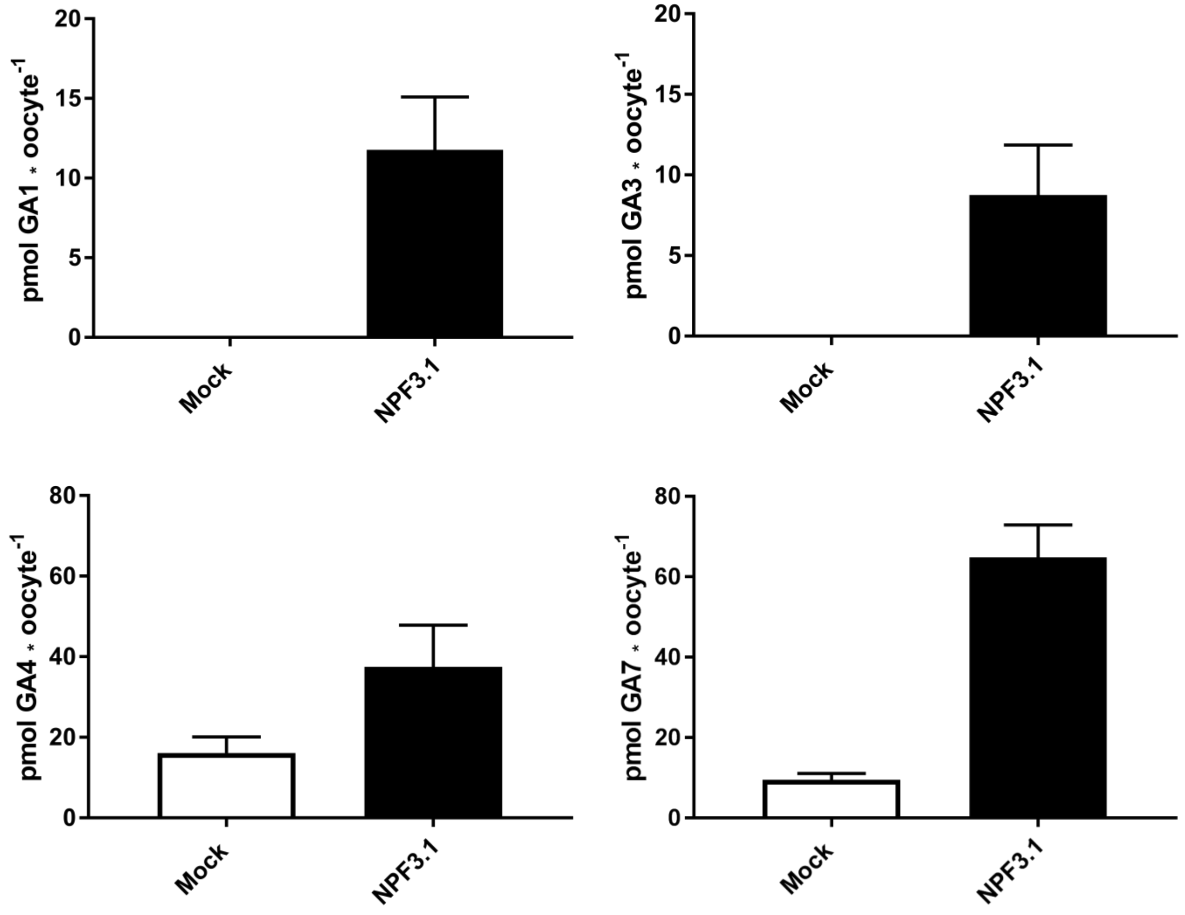

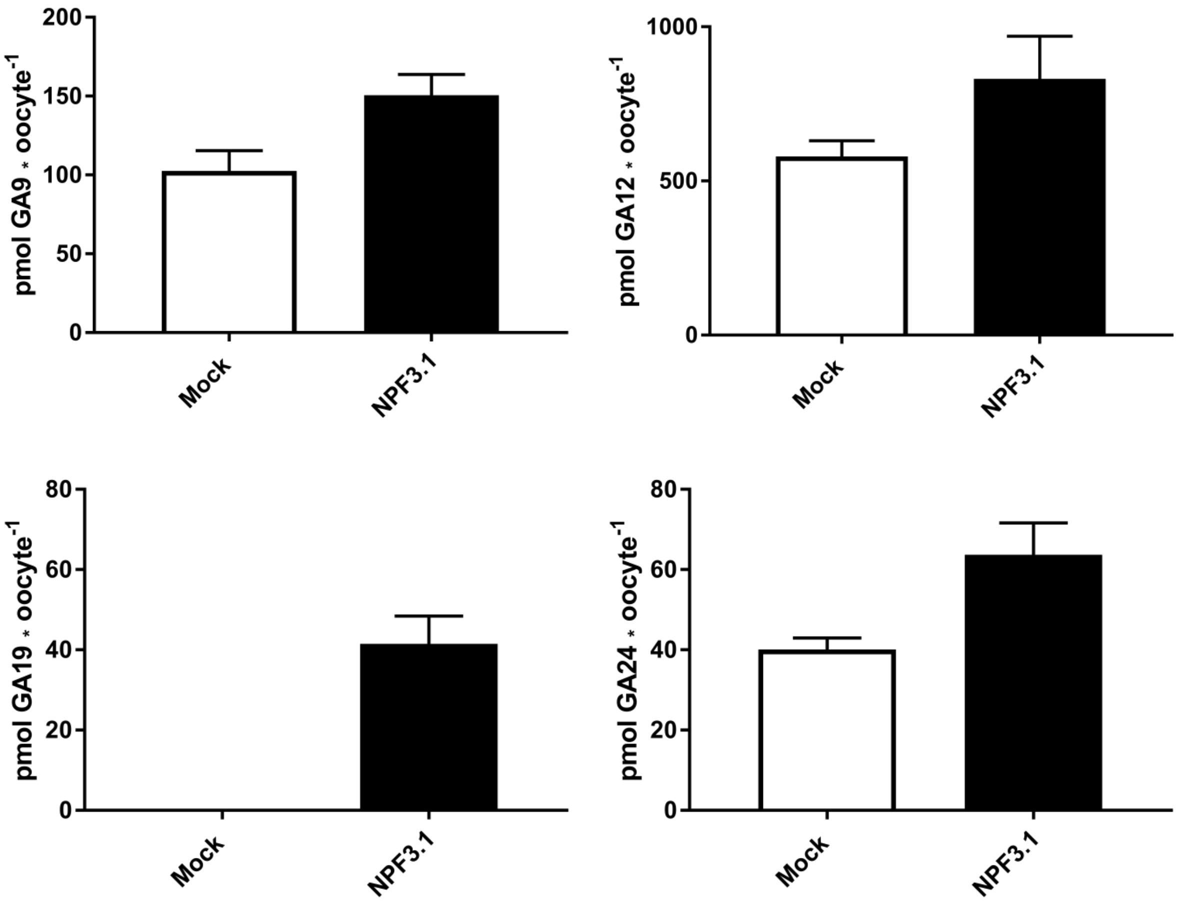


**Supplementary Figure 1.** Expanded list of NPF3.1 GA substrates. Membrane non-permeable GAs (GA1, GA3 and GA19) were assayed in MES based kulori pH 5 for 1 h. Moderately membrane permeable GAs (GA4, GA7 and GA24) were assayed in MES based kulori pH 5.5 for 1 h. Highly membrane permeable GAs (GA9 and GA12) were assayed in MES based kulori pH 6 for 1 h. All transport events are statistically significant (Holm Sidak one-way ANOVA p = 0.05).

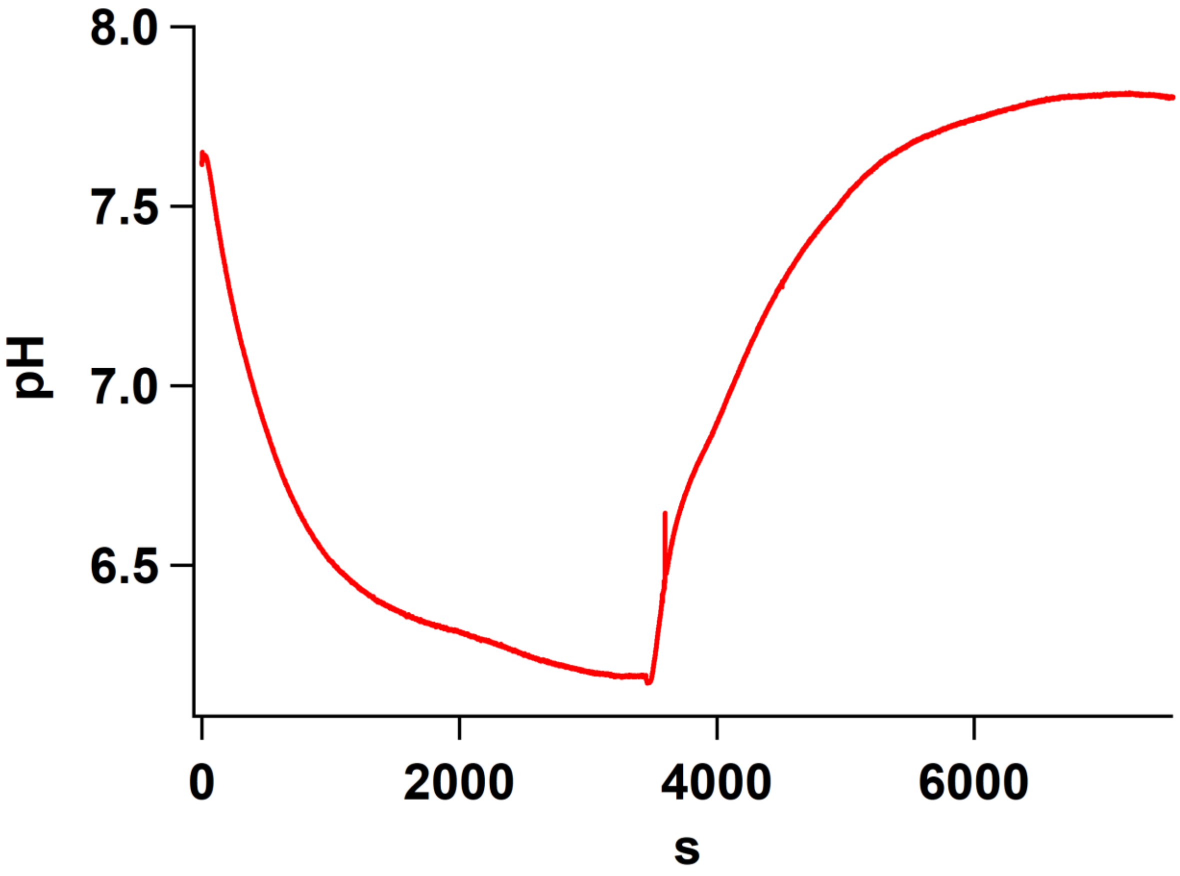


**Supplementary Figure 2.** Cytosolic pH of NPF7.3 expressing oocytes upon exposure to pH 5.0 for 60 mins followed by 60 mins in pH 7.4

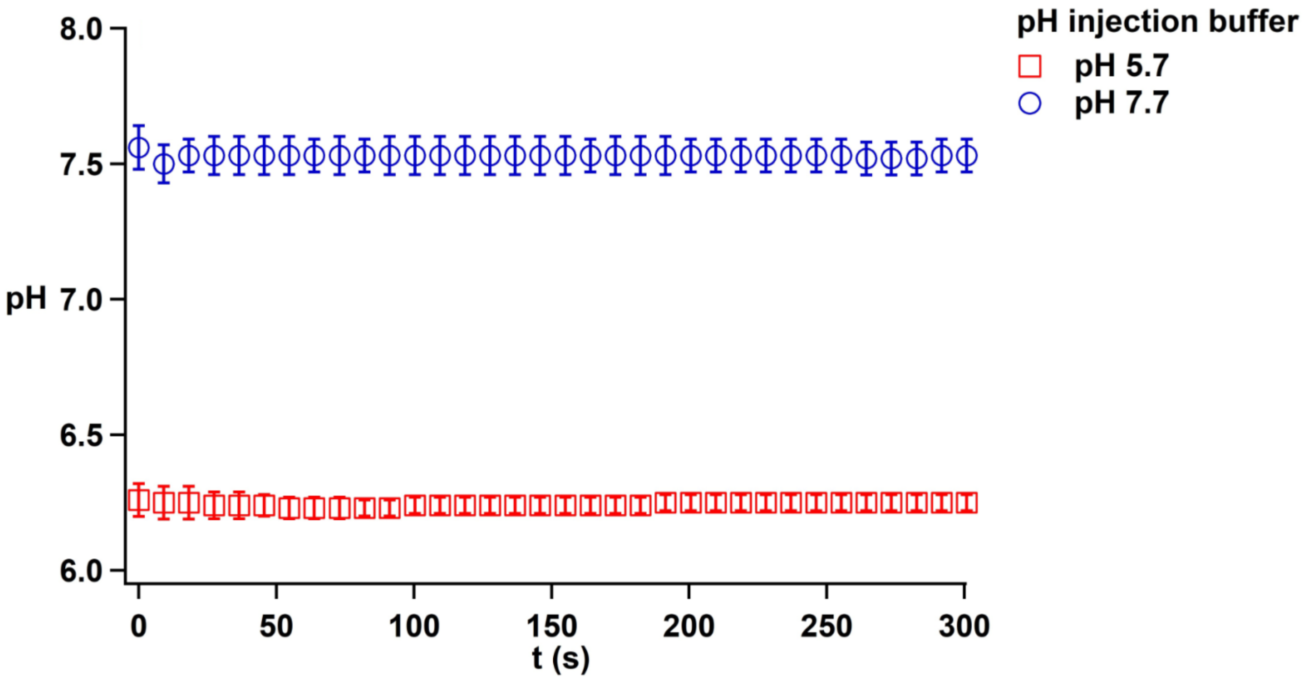


**Supplementary Figure 3.** Internal pH of oocytes injected with 50.6 nl 0.5 M TRIS 50 mM EGTA adjusted to pH 7.7 with 0.5 M MES (blue circles) and internal pH of oocytes injected with 50.6 nl 0.5 M MES 50 mM EGTA adjusted to 5.7 with 0.5 M TRIS (red squares).

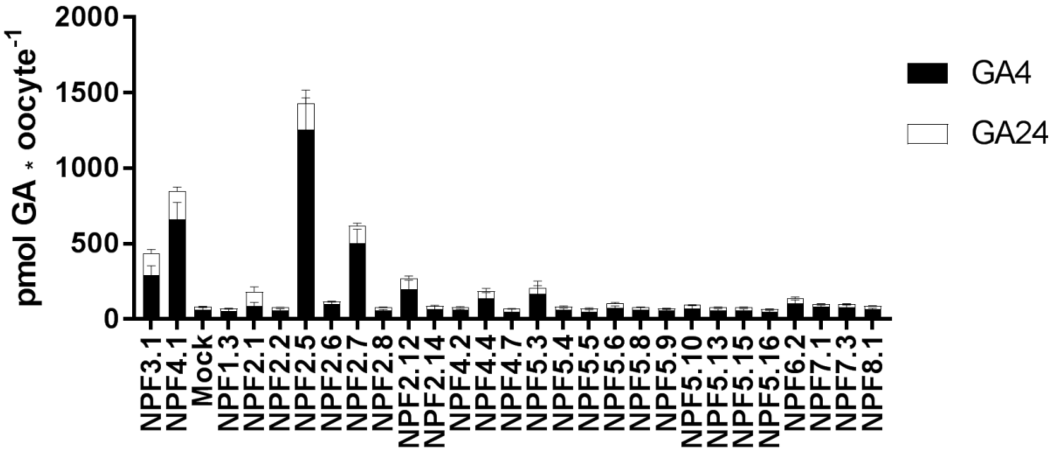
pmol GA * oocyte-1
2000
1500
1000
500
0
GA4
GA24
NPF3.1
NPF4.1
Mock
NPF1.3
NPF2.1
NPF2.2
NPF2.5
NPF2.6
NPF2.7
NPF2.8
NPF2.12
NPF2.14
NPF4.2
NPF4.4
NPF4.7
NPF5.3
NPF5.4
NPF5.5
NPF5.6
NPF5.8
NPF5.9
NPF5.10
NPF5.13
NPF5.15
NPF5.16
NPF6.2
NPF7.1
NPF7.3
NPF8.1

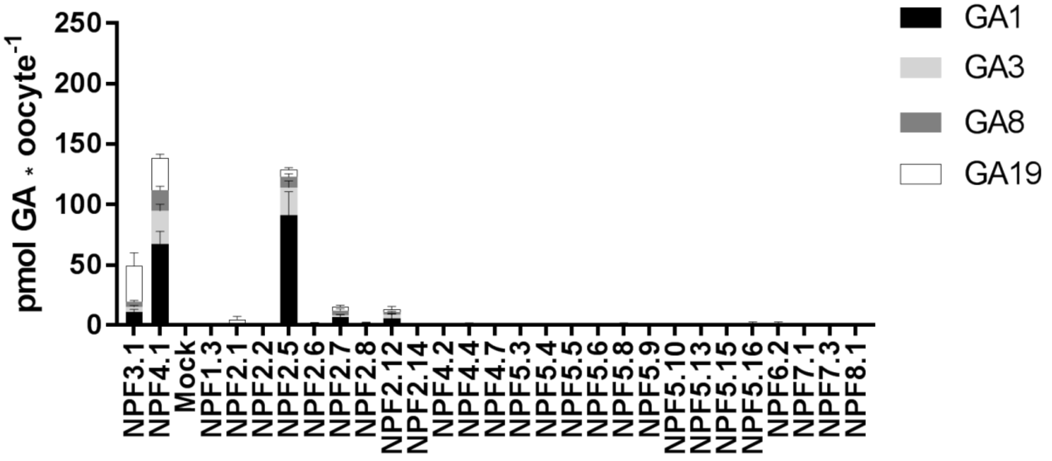
pmol GA * oocyte-1
250
200
150
100
50
0
GA1
GA3
GA8
GA19
NPF3.1
NPF4.1
Mock
NPF1.3
NPF2.1
NPF2.2
NPF2.5
NPF2.6
NPF2.7
NPF2.8
NPF2.12
NPF2.14
NPF4.2
NPF4.4
NPF4.7
NPF5.3
NPF5.4
NPF5.5
NPF5.6
NPF5.8
NPF5.9
NPF5.10
NPF5.13
NPF5.15
NPF5.16
NPF6.2
NPF7.1
NPF7.3
NPF8.1

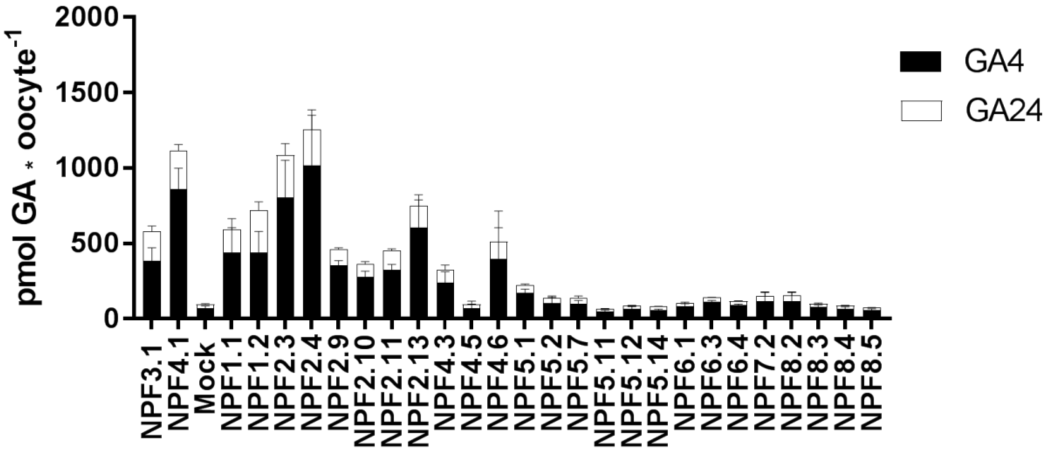


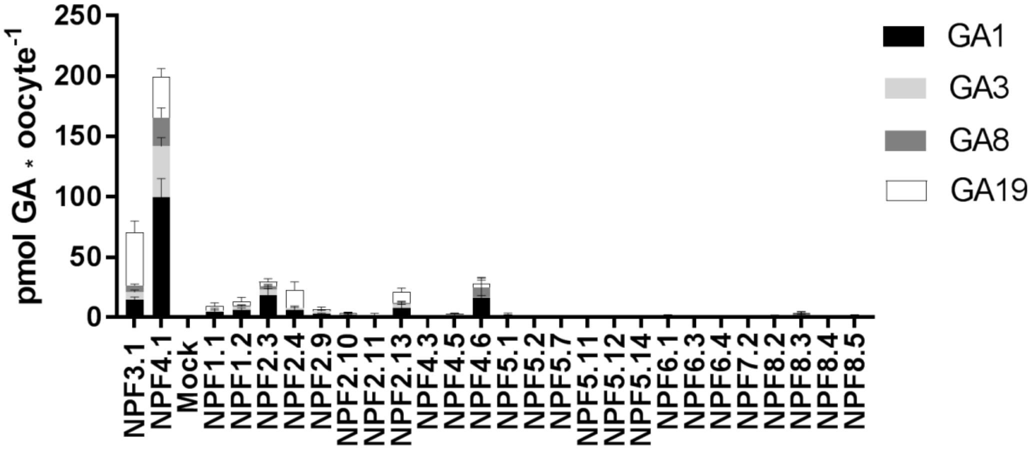


**Supplementary Figure 4.** Un-normalized data of first and second half of the quantitative screen. Both assays included NPF3.1, NPF4.1 and Mock in order to normalize. Oocytes (n = 5-6) were exposed to a mix of 50 µM GA1, 50 µM GA3, 100 µM GA4, 50 µM GA8, 50 µM GA19 and 50 µM GA24 in pH kulori 5.5 for 1h. Oocyte GA content was analyzed using LC-MS. For ease of visualization, data is divided to show accumulation of either membrane permeable or membrane non-permeable GAs.

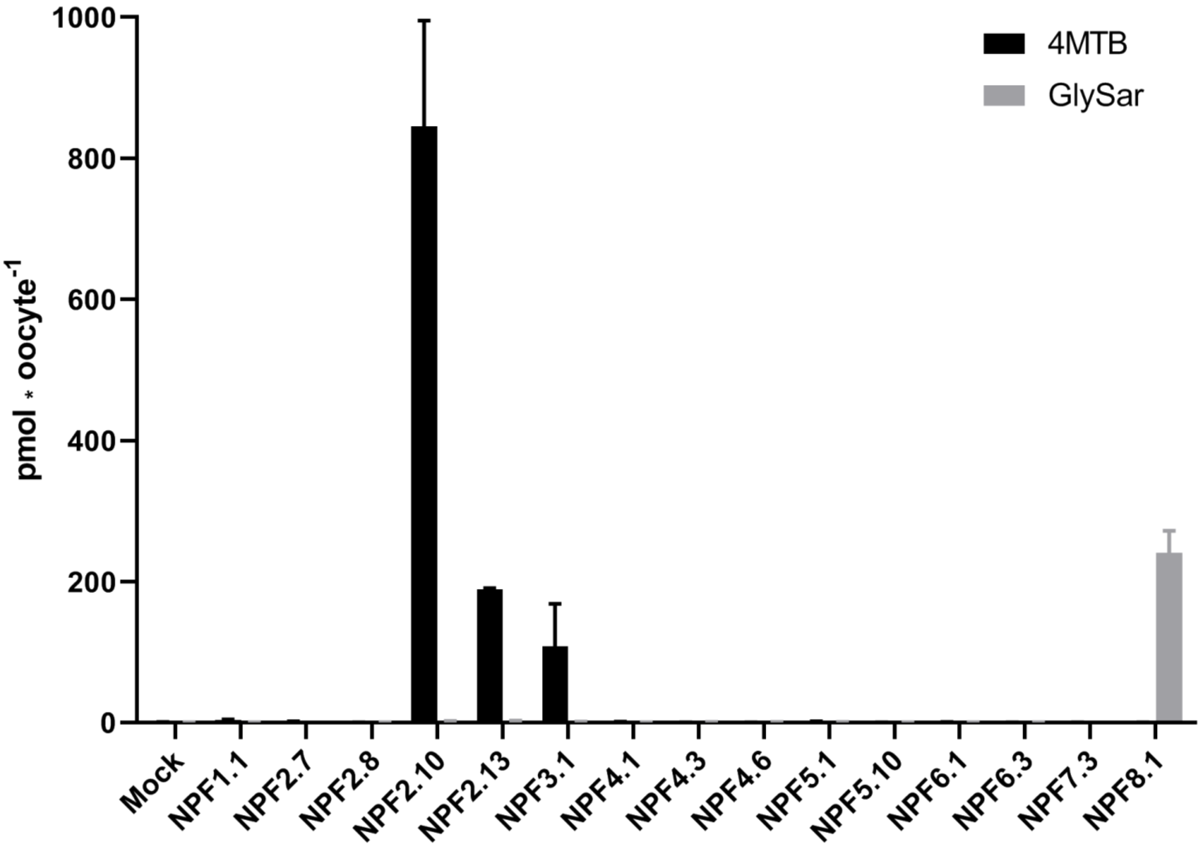


**Supplementary Figure 5.** Aliphatic glucosinolate and peptides transport are confined to branches in the NPF family. Chosen Arabidopsis NPF members were subjected to a mix of 500 µM 4-methylthio-3-butenyl aliphatic glucosinolate and 500 µM un-metabolizable glycyl-sarcosine dipeptide in MES based kulori pH 5 for 1 h and analyzed on LC-MS/MS.

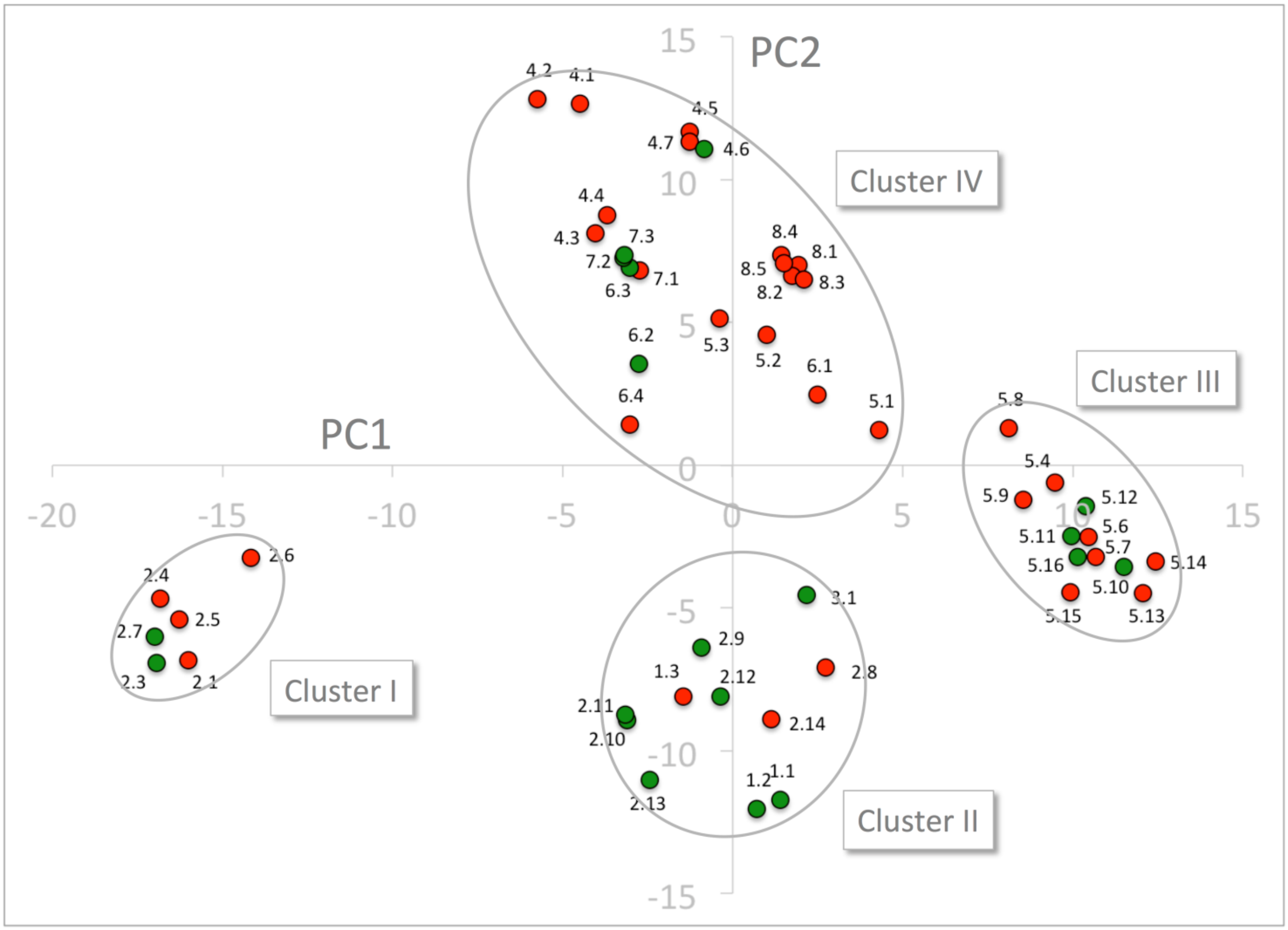


**Supplementary Figure 6.** Projection of nitrate transporting NPFs onto Principal Component Analysis of the 51 NPF sequences expressed by z-scales of the 51 cavity residues. Nitrate transporting transporters are shown as green dots and non-nitrate transporting transporters as red dots. The four clusters are marked by ellipses. PC1 and PC2 refer to the first and second principal components, respectively.

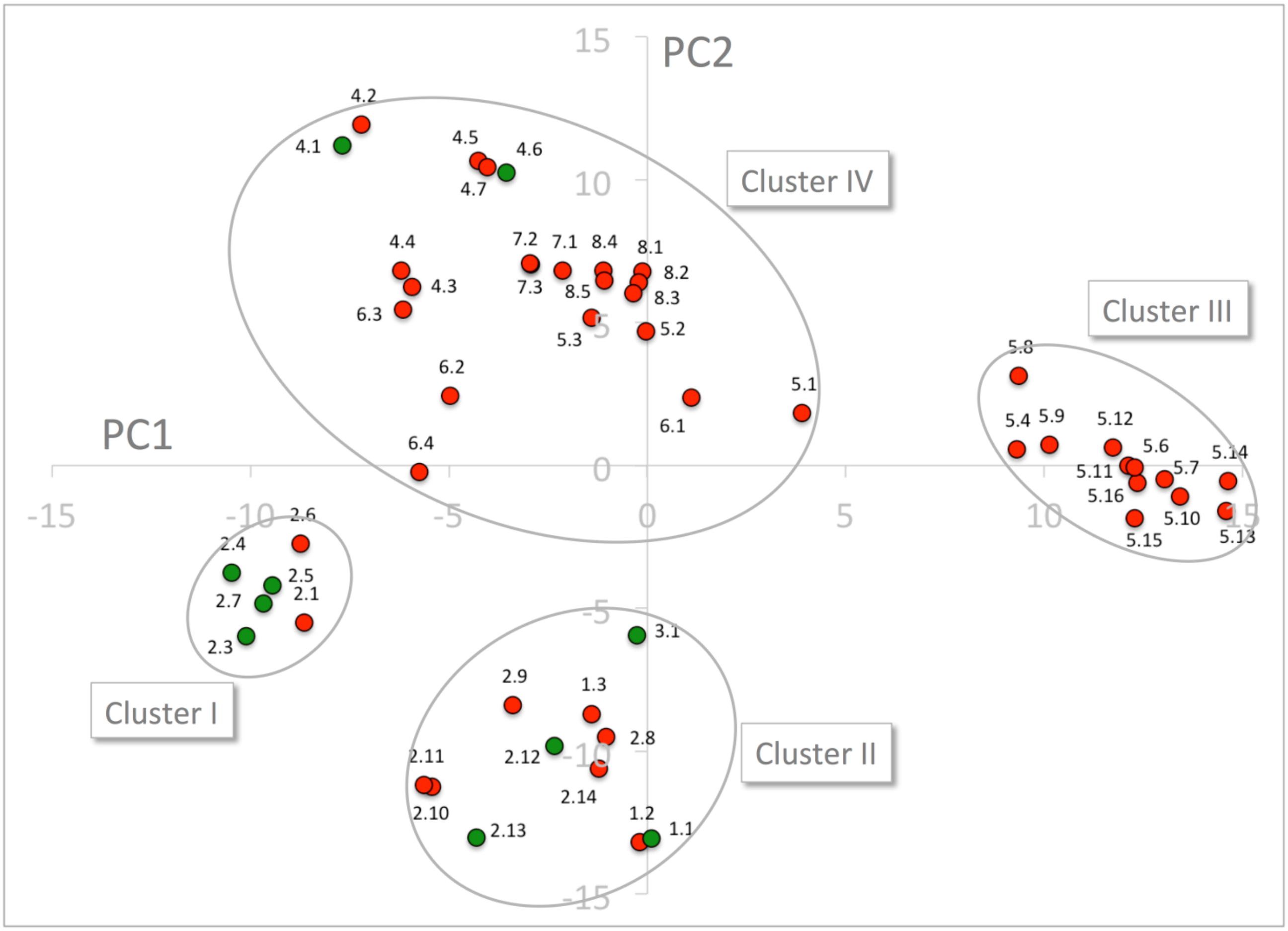


**Supplementary Figure 7.** Principal Component Analysis of the 51 NPF sequences expressed by z-scales of the 48 cavity residues that excludes the ExxE[K/R] motif. GA transporting and GA non-transporting transporters are shown as green and red dots, respectively. The four clusters are marked by ellipses. PC1 and PC2 refer to the first and second principal components, respectively.

| Gene | Substrates | Non-substrate GAs | Ref |
|---|---|---|---|
| AtNPF1.1 | GA1, GA3, GA4, ABA, JA-Ile | - | Chiba et al. 2015 |
| | Nitrate | - | Hsu et al. 2013 |
| AtNPF1.2 | GA1, GA3, GA4, JA-Ile | - | Chiba et al. 2015 |
| | Nitrate | - | Hsu et al. 2013. |
| AtNPF2.3 | GA1, GA3, (GA4) | - | Chiba et al. 2015 |
| | Nitrate | - | Taochy et al. 2015 |
| AtNPF2.4 | GA1, GA3, (GA4) | - | Chiba et al. 2015 |
| | Chloride | - | Li et al. 2016[1] |
| AtNPF2.5 | GA1, GA3, (GA4), ABA | - | Chiba et al. 2015 |
| | GA3 | - | Kanno et al. 2016 |
| | Chloride | - | Li et al. 2016[2] |
| AtNPF2.6 | (GA1, GA4, JA-Ile) | GA3 | Chiba et al. 2015 |
| AtNPF2.7 | GA1, GA3 | GA4 | Chiba et al. 2015 |
| | Nitrate | | Segonzac et al. 2007 |
| AtNPF2.10 | **GA1, GA3** | GA4 | Chiba et al. 2015 |
| | **GA3,** ABA, JA, JA-Ile, TA | **GA1,** GA4, GA8, GA9, GA20 | Saito et al. 2015 |
| | **GA3** | - | Kanno et al. 2016 |
| | - | **GA3,** GA4 | Tal et al. 2016 |
| | 4MTB | - | Nour-Eldin et al. 2012 |
| | I3M | - | Jorgensen et al. 2017 |
| AtNPF2.11 | GA3 | - | Saito et al. 2015 |
| | GA3 | - | Tal et al. 2016 |
| | 4MTB | - | Nour-Eldin et al. 2012 |
| | I3M | - | Jorgensen et al. 2017 |
| AtNPF2.12 | GA1, GA3 | GA4 | Chiba et al. 2015 |
| | Nitrate | - | Almagro et al. 2008 |
| AtNPF2.13 | GA1, GA3, (GA4) | - | Chiba et al. 2015 |
| | Nitrate | - | Fan et al. 2009 |
| | 4MTB | - | Nour-Eldin et al. 2012 |
| | I3M | - | Jorgensen et al. 2017 |
| AtNPF3.1 | GA1, GA3 | **GA4** | Chiba et al. 2015 |
| | GA1, GA3, **GA4**, GA8, **GA20** | GA9, GA12 | Tal et al. 2016 |
| | GA1, GA3, **GA4**, GA8, GA19 | GA9, GA12, GA15, **GA20**, GA24, | David et al. 2016 |
| | - | GA44, GA53 | David et al. 2016 |
| | Nitrate, Nitrite | - | Pike et al. 2014 |
| AtNPF4.1 | GA3, ABA, JA | - | Kanno et al. 2012 |
| | GA1, GA3, GA4 | - | Chiba et al. 2015 |
| | GA3 | - | Saito et al. 2015 |
| | GA3, GA4, GA8, GA20 | GA9, GA12 | Tal et al. 2016 |
| AtNPF4.2 | GA1, GA3, (GA4) | - | Chiba et al. 2015 |
| | ABA | - | Kanno et al. 2012 |
| AtNPF4.5 | (GA1, GA3), ABA | - | Chiba et al. 2015 |
| | ABA | - | Kanno et al. 2012 |
| AtNPF4.6 | (GA1, GA3), ABA | - | Chiba et al. 2015 |
| | ABA | - | Kanno et al. 2012 |
| | Nitrate | - | Huang et al. 1999 |
| AtNPF5.1 | GA1, GA3, (GA4) | - | Chiba et al. 2015 |
| AtNPF5.2 | GA1, GA3, (GA4), ABA | - | Chiba et al. 2015 |
| | Peptides | - | Karim et al. 2007 |
| AtNPF5.3 | (GA1, GA3, ABA) | GA4 | Chiba et al. 2015 |
| AtNPF5.6 | (GA1, GA3, GA4) | - | Chiba et al. 2015 |
| AtNPF5.7 | GA1, **GA3, GA4**, JA-Ile | - | Chiba et al. 2015 |
| | - | **GA3, GA4** | Tal et al. 2016 |
| AtNPF5.8 | (GA1, GA4) | GA3 | Chiba et al. 2015 |
| AtNPF5.9 | (GA1, GA4) | GA3 | Chiba et al. 2015 |
| AtNPF6.3 | (GA1, GA4) | GA3 | Chiba et al. 2015 |
| | Nitrate | - | Tsay et al. 1993 |
| | IAA | - | Krouk et al. 2010 |
| AtNPF8.2 | (GA1, GA4, ABA), JA-Ile | GA3 | Chiba et al. 2015 |
| | Peptides | - | Komarova et al. 2008 |
| | Peptides | - | Hammes et al. 2010 |
| AtSWEET13 | GA1, GA3, GA4, (GA9), GA12, GA15, | GA8 | Kanno et al. 2016 |
| | GA19, GA20 GA24, GA44, GA53 | - | Kanno et al. 2016 |

| | | | |
|---|---|---|---|
| | Sucrose | - | Chen et al. 2012 |
| AtSWEET14 | GA1, GA3, GA4, GA8, GA9, GA12, | - | Kanno et al. 2016 |
| | GA15, GA19, GA20 GA24, GA44, | - | Kanno et al. 2016 |
| | GA53 | - | Kanno et al. 2016 |
| | Sucrose | | Chen et al. 2012 |

**Supplementary Table 1.** Summary of substrates of published GA transporters. Lack of consensus between GA transport assays is highlighted in **Bold**. Data presented in parentheses are regarded as positive transport events, but uncertainty exists. (Tsay, Schroeder et al. 1993, Huang, Liu et al. 1999, Karim, Holmstrom et al. 2007, Segonzac, Boyer et al. 2007, Almagro, Lin et al. 2008, Komarova, Thor et al. 2008, Fan, Lin et al. 2009, Hammes, Meier et al. 2010, Krouk, Lacombe et al. 2010, Chen, Qu et al. 2012, Kanno, Hanada et al. 2012, Nour-Eldin, Andersen et al. 2012, Hsu and Tsay 2013, Pike, Gao et al. 2014, Chiba, Shimizu et al. 2015, Saito, Oikawa et al. 2015, Taochy, Gaillard et al. 2015, David, Berquin et al. 2016, Kanno, Oikawa et al. 2016, Li, Byrt et al. 2016, Li, Qiu et al. 2016, Tal, Zhang et al. 2016, Jørgensen, Xu et al. 2017)

| Gene | AGI | Direction | Primer sequence | CDS Origin |
|---|---|---|---|---|
| NPF1.1 | At3g16180 | Forward<br>Reverse | GGCTTAAUATGGAGAACCCTCCCAATGA<br>GGTTTAAUTTAATTGGTTTTAACAACTGGACTTAGATCTTC | Courtesy of Wolf B. Frommer |
| NPF1.2 | At1g52190 | - | - | uNCDF |
| NPF1.3 | At5g11570 | - | - | uNCDF |
| NPF2.1 | At3g45720 | Forward<br>Reverse | GGCTTAAUATGGCTGGTTTAGTATTATCTA<br>GGTTTAAUTTAGTTTGTAACATCTTTAGGA | Cloned from cDNA |
| NPF2.2 | At3g45690 | - | - | uNCDF |
| NPF2.3 | At3g45680 | - | - | uNCDF |
| NPF2.4 | At3g45700 | Forward<br>Reverse | GGCTTAAUATGGCTAATTCAGACTCTGGTGACAAAGAA<br>GGTTTAAUCTAGTTTGTAACATCTTTAAGATCTTGTTCATGATC | Courtesy of Wolf B. Frommer |
| NPF2.5 | At3g45710 | Forward<br>Reverse | GGCTTAAUATGGCTGATTCAAAATCTGGTGAC<br>GGTTTAAUCTAGGTTTTAACATCTTTAGGATCTTGTTCATG | Cloned from cDNA |
| NPF2.6 | At3g45660 | - | - | uNCDF |
| NPF2.7 | At3g45650 | Forward<br>Reverse | GGCTTAAUATGGCTAGTTCAGTTACTG<br>GGTTTAAUTCAGTGAGAGACATTTGC | Cloned from cDNA |
| NPF2.8 | At5g28470 | - | - | uNCDF |
| NPF2.9 | At1g18880 | Forward<br>Reverse | GGCTTAAUATGGAGGTTGAGAAGACAG<br>GGTTTAAUTTACACTGACACCTTATCAAAC | Courtesy of Wolf B. Frommer |
| NPF2.10 | At3g47960 | Forward<br>Reverse | GGCTTAAUATGAAGAGCAGAGTCATT<br>GGTTTAAUTCAGACAGAGTTCTTGTC | RIKEN BRC |
| NPF2.11 | At5g62680 | Forward<br>Reverse | GGCTTAAUATGGAGAGAAAGCCTCTTGAAC<br>GGTTTAAUTCAGGCAACGTTCTTGTCTTG | Nour Eldin et al. 2012) |
| NPF2.12 | At1g27080 | Forward<br>Reverse | GGCTTAAUATGGGAGTTGTTGAGAATCGG<br>GGTTTAAUCTAACTTGGAGATTGATCGTGTCTC | Courtesy of Eilon Shani |
| NPF2.13 | At1g69870 | Forward<br>Reverse | GGCTTAAUATGGTTTTGGAGGATAGAAAGGAC<br>GGTTTAAUTCATTTCATCGATTTCTTCGAAGTCATCTC | Nour Eldin et al. 2012) |
| NPF2.14 | At1g69860 | Forward<br>Reverse | GGCTTAAUATGGACAATGAGAAAGGGACAAG<br>GGTTTAAUTTATTGTTCATTACTATAGTTTCTGTAACGATACC | (Nour Eldin et al. 2012) |
| NPF3.1 | At1g68570 | Forward<br>Reverse | GGCTTAAUATGGAGGAGCAAAGCAAGAA<br>GGTTTAAUTCATTCATCAACTAAACTCCTA | RIKEN BRC |
| NPF4.1 | At3g25260 | Forward<br>Reverse | GGCTTAAUATGCAGATTGAGATGGAAGAG<br>GGTTTAAUCTAATATCTTTTCGCCCAG | Cloned from cDNA |
| NPF4.2 | At3g25280 | - | - | uNCDF |
| NPF4.3 | At1g59740 | Forward<br>Reverse | GGCTTAAUATGGCAGAGATAAACAAACAAAGCA<br>GGTTTAAUCTAAATGTTCTCATCACCCACAAC | RIKEN BRC |
| NPF4.4 | At1g33440 | - | - | uNCDF |
| NPF4.5 | At1g27040 | Forward<br>Reverse | GGCTTAAUATGGAAGTAGAAATGCATGGTGA<br>GGTTTAAUTCAACTTATTGAACCAGTTGAGATATAC | RIKEN BRC |
| NPF4.6 | At1g69850 | Forward<br>Reverse | GGCTTAAUATGGAAGTGGAAGAAGAGGTCTC<br>GGTTTAAUTTAGCTTCTTGAACCAGTTGATCTATAC | Courtesy of Wolf B. Frommer |
| NPF4.7 | At5g62730 | - | - | uNCDF |
| NPF5.1 | At2g40460 | Forward<br>Reverse | GGCTTAAUATGGAGGCTGCAAAAGTTTACAC<br>GGTTTAAUTTAGATACTAAGAGGAGATGTGTCTAAGGC | Courtesy of Wolf B. Frommer |
| NPF5.2 | At5g46050 | Forward<br>Reverse | GGCTTAAUATGACAGTAGAAGAGGTAGGAGAC<br>GGTTTAAUTTATTCAGTCTCTTTCATTTCCACCTC | RIKEN BRC |
| NPF5.3 | At5g46040 | Forward<br>Reverse | GGCTTAAUATGACAGTAGAAGAGGTAGG<br>GGTTTAAUTTACTCATTGTAGTTATCTACC | Cloned from cDNA |
| NPF5.4 | At3g54450 | - | - | uNCDF |
| NPF5.5 | At2g38100 | - | - | uNCDF |
| NPF5.6 | At2g37900 | - | - | uNCDF |
| NPF5.7 | At3g53960 | - | - | uNCDF |
| NPF5.8 | At5g14940 | - | - | uNCDF |
| NPF5.9 | At3g01350 | - | - | uNCDF |
| NPF5.10 | At1g22540 | Forward<br>Reverse | GGCTTAAUATGTCGATCTCCGGCGCT<br>GGTTTAAUTTAACTGGTGTCGAGCCTTT | Cloned from cDNA |
| NPF5.11 | At1g72130 | Forward<br>Reverse | GGCTTAAUATGGCTATCACCTACTCCTCC<br>GGTTTAAUTTAAAAGGTGTTTGATCTGCTGTAGACAT | RIKEN BRC |
| NPF5.12 | At1g72140 | Forward<br>Reverse | GGCTTAAUATGTCGACATCCATCGGCG<br>GGTTTAAUCTACTTTGGGCTGTTGTAGAGATAG | RIKEN BRC |

| | | | | |
|---|---|---|---|---|
| NPF5.13 | At1g72125 | Forward<br>Reverse | GGCTTAAUATGACGACGACTTCCAAAAC<br>GGTTTAAUCTACACTACGTCCACCCG | Cloned from cDNA |
| NPF5.14 | At1g72120 | Forward<br>Reverse | GGCTTAAUATGACGACTACTTCAGAAATTTCTCT<br>GGTTTAAUCTACACTCGATCCACTCGACG | RIKEN BRC |
| NPF5.15 | At1g22570 | Forward<br>Reverse | GGCTTAAUATGAAGATACCAGAGGAAGAAGTTGC<br>GGTTTAAUTTAGACTTGGTCTAGCCGACG | Cloned from cDNA |
| NPF5.16 | At1g22550 | Forward<br>Reverse | G GGCTTAAUATGGCGATAGCCGAAGAAGAA<br>GGTTTAAUTTAGACTTGATCTACACGGCG | Courtesy of Wolf B. Frommer |
| NPF6.1 | At5g13400 | Forward<br>Reverse | GGCTTAAUATGGTTGCTTCTGAGATTAAATCCC<br>GGTTTAAUTTAAAGGACAGCACTACTCTTGTCTTCC | Courtesy of Wolf B. Frommer |
| NPF6.2 | At2g26690 | Forward<br>Reverse | GGCTTAAUATGGAGAGCAAAGGGAGTTGG<br>GGTTTAAUTCAGCAGTCTTCAACTGAAAATCC | RIKEN BRC |
| NPF6.3 | At1g12110 | Forward<br>Reverse | GGCTTAAUATGTCTCTTCCTGAAACTAAATCTG<br>GGTTTAAUATGACCCATTGGAATACTCG | Courtesy of Wolf B. Frommer |
| NPF6.4 | At3g21670 | Forward<br>Reverse | GGCTTAAUATGGTTCATGTGTCATCATCTCATG<br>GGTTTAAUTCAAGGAATGTCCTTAAGCTCAA | Courtesy of Wolf B. Frommer |
| NPF7.1 | At5g19640 | Forward<br>Reverse | GGCTTAAUATGGCCGCTATGGATCCG<br>GGTTTAAUAGTTTGAACAAGGTTGAGTCTTT | Cloned from cDNA |
| NPF7.2 | At4g21680 | Forward<br>Reverse | GGCTTAAUATGGATCAAAAAGTTAGACAGT<br>GGTTTAAUTCAGACTTCCTCCTCTTCAGTTA | Courtesy of Wolf B. Frommer |
| NPF7.3 | At1g32450 | Forward<br>Reverse | GGCTTAAUATGTCTTGCCTAGAGATTTAT<br>GGTTTAAUTTAGACTTTAGAATCCTTCTCT | Courtesy of Wolf B. Frommer |
| NPF8.1 | At3g54140 | Forward<br>Reverse | GGCTTAAUATGGAAGAAAAAGATGTGTATACGC<br>GGTTTAAUTCAATGTGCTCGACCAACAGC | Courtesy of Wolf B. Frommer |
| NPF8.2 | At5g01180 | Forward<br>Reverse | GGCTTAAUATGGAAGATGACAAGGATATATACACAAA<br>GGTTTAAUTCAAAGCGCATGCCCGGT | Courtesy of Wolf B. Frommer |
| NPF8.3 | At2g02040 | - | - | uNCDF |
| NPF8.4 | At2g02020 | - | - | uNCDF |
| NPF8.5 | At1g62200 | Forward<br>Reverse | GGCTTAAUATGGTGAATTCGAATGAAGAAGACG<br>GGTTTAAUTTACAAAGCCTTCTTCTTTGTGTGC | Courtesy of Wolf B. Frommer |

**Supplementary Table 2.** List of primers and CDS origin. Uracil containing Non-clonal DNA fragments (uNCDF).

| Analyte | RT [min] | Q1 [*m/z*] | Q3 [*m/z*] | CE [eV] |
|---|---|---|---|---|
| GA1 [M-H]$^{-}$ | 2.29 | 347.16 | 259.1$^{Q}$ | 16 |
| | | 347.16 | 145.2 | 28 |
| GA3 [M-H]$^{-}$ | 2.27 | 345.2 | 243.1$^{Q}$ | 25 |
| | | 345.2 | 239.1 | 12 |
| | | 345.2 | 221.1 | 21 |
| GA4 [M-H]$^{-}$ | 2.62 | 331.1 | 243.0$^{Q}$ | 16 |
| | | 331.1 | 225.1 | 15 |
| | | 331.1 | 213.0 | 30 |
| | | 331.1 | 287,1 | 19 |
| GA7 [M-H]$^{-}$ | 2.64 | 329.15 | 223.1$^{Q}$ | 16 |
| | | 329.15 | 211.1 | 21 |
| | | 329.15 | 155.0 | 27 |
| GA8 [M-H]$^{-}$ | 2.05 | 363.2 | 275.1$^{Q}$ | 15 |
| | | 363.2 | 257.1 | 15 |
| | | 363.2 | 119.2 | 23 |
| | | 363.2 | 319,1 | 11 |
| GA9 [M-H]$^{-}$ | 2.82 | 315.2 | 271.1$^{Q}$ | 16 |
| | | 315.2 | 253.1 | 23 |
| GA12 [M-H]$^{-}$ | 2.89 | 331.2 | 313.1$^{Q}$ | 24 |
| | | 331.2 | 287.1 | 22 |
| GA19 [M-H]$^{-}$ | 2.38 | 331.2 | 273.0$^{Q}$ | 24 |
| | | 331.2 | 203.0 | 29 |
| | | 331.2 | 317.0 | 22 |
| | | 331.2 | 229.0 | 29 |
| GA24 [M-H]$^{-}$ | 2.61 | 345.1 | 257.0$^{Q}$ | 25 |
| | | 345.1 | 213.0 | 25 |
| | | 345.1 | 301.0 | 16 |
| 4-methylthio-3-butenyl [M-H]$^{-}$ | 2.27 | 420.0 | 97.0$^{Q}$ | 23 |
| | | 420.0 | 259.0 | 23 |
| | | 420.0 | 75.0 | 30 |
| Glycyl-sarcosine [M+H]$^{+}$ | 0.58 | 147.1 | 90.2$^{Q}$ | -8 |
| | | 147.1 | 44.4 | -17 |
| JA [M-H]$^{-}$ | 2.45 | 209.1 | 59.3$^{Q}$ | 11 |
| JA-Ile [M-H]$^{-}$ | 2.49 | 322.0 | 130.1$^{Q}$ | 17 |

| | | | | |
|---|---|---|---|---|
| ABA $[M-H]^-$ | 2.34 | 263.0 | 153.1$^{Q}$ | 7 |
| | | 263.0 | 151.0 | 7 |
| OPDA $[M-H]^-$ | 2.79 | 291.0 | 165.1$^{Q}$ | 17 |
| | | 291.0 | 273.1 | 13 |
| Sinigrin $[M-H]^-$ (IS) | 1.60 | 358.0 | 97.0$^{Q}$ | 22 |
| | | 358.0 | 79.0 | 30 |
| | | 358.0 | 259.0 | 20 |

**Supplementary Table 3.** Multiple reaction monitoring transitions for analyte detection by LC-MS/MS. $^{Q}$Quantifier ion used for quantification of the respective analyte. Other transitions were used for compound identification together with retention times compared to those of standards. IS = internal standard.
